# Lactate production from lactose-rich wastewater: A comparative study on reactor configurations to maximize conversion rates and efficiencies

**DOI:** 10.1101/2024.10.16.618679

**Authors:** Monika Temovska, Richard Hegner, Andrés E. Ortiz-Ardila, Joseph G. Usack, Largus T. Angenent

**Affiliations:** Environmental Biotechnology Group, Department of Geosciences, University of Tübingen, Schnarrenbergstr. 94-96, 72076 Tübingen, Germany; AG Angenent, Max Planck Institute for Biology, Max Planck Ring 5, 72076 Tübingen, Germany; Department of Food Science and Technology, University of Georgia, Athens, Georgia; Department of Biological and Chemical Engineering, Aarhus University, Gustav Wieds vej 10D, 8000 Aarhus C, Denmark; The Novo Nordisk Foundation CO_2_ Research Center (CORC), Aarhus University, Gustav Wieds vej 10C, 8000 Aarhus C, Denmark; Cluster of Excellence – Controlling Microbes to Fight Infections, University of Tübingen, Auf der Morgenstelle 28, 72074 Tübingen, Germany

**Keywords:** Sugar-rich wastewater, lactose, acid whey, lactate, D-lactate, thermophilic fermentation, reactor microbiome, *Lactobacillus* spp

## Abstract

About 90% of global lactate production is derived from bacterial fermentation of sugars *via* pure cultures of homofermentative bacteria in batch mode. Acid whey, which is a wastewater from the yogurt industry with lactose and galactose as the main sugars, can be used as an alternative substrate for the commercial production of lactate. Operating open cultures of microbial consortia (*i.e*., reactor microbiomes) reduces the costs of lactate production by circumventing sterilization, while continuous operation achieves higher productivity at shorter production times. Homofermentation can be achieved by maintaining acidic and thermophilic conditions, while product formation in continuous systems can be increased with biomass retention strategies. To find the best reactor configuration for lactate production from acid whey, we operated three different reactor configurations: **(1)** an upflow anaerobic sludge blanket (UASB) reactor; **(2)** an anaerobic filter reactor (AFR); and **(3)** an anaerobic continuously stirred tank reactor (CSTR) with a hollow-fiber membrane module. We operated at different hydraulic retention times (HRTs) to find the optimum parameters to maximize the lactose and galactose-into-lactate (LG-into-LA) conversion efficiency. We did not use an inoculum but enriched the endogenous D-lactate-producing *Lactobacillus* spp. that later dominated the reactor microbiomes (> 90% relative abundance). Undissociated lactic acid concentrations of more than 60 mmol C L^-1^ inhibited the microbiomes. We alleviated the inhibition effect by shortening the HRT to 0.6 days and using diluted acid-whey substrate (1.67-fold dilution) to achieve almost complete conversion of the acid-whey sugars to lactate. At the 0.6-day HRT commencement, the AFR and CSTR performed better than the UASB reactor due to their better cell retention abilities. During the period between Day 365-384, we experienced an error in the pH control of the CSTR system during which the pH value dropped to 4.3. After this pH-error period, the LG-into-LA conversion efficiency for the CSTR considerably improved and surpassed the AFR. We achieved the highest lactate conversion rate of 1256 ± 46.3 mmol C L^-1^ d^-1^ (1.57 ± 0.06 g L^-1^ h^-1^) at a LG-into-LA conversion efficiency of 82.2 ± 3.4% (in mmol C), with a yield of 0.85 ± 0.02 mmol C mmol C^-1^ (product per consumed substrate) for the CSTR.

## 1. Introduction

Lactate is a versatile platform chemical with a broad range of applications across the food, pharmaceutical, cosmetic, detergent, and dairy industries. There has been a recent expansion in the market for lactate, particularly for the production of polylactate (PLA) (López-Gómez et al., 2019). Furthermore, acid whey can be used as a complex substrate within the carboxylate platform, with lactate being an intermediate product, to produce the higher-value medium-chain carboxylates (MCCs), which are bio-fuel precursors (Xu et al., 2018). Throughout this text, we use the term carboxylates to refer to mixtures of both dissociated carboxylates and their corresponding undissociated carboxylic acids. Chemically, lactate is produced *via* the hydrolysis of lactonitrile, which is a process that yields a racemic mixture of D- and L-lactate. Compared to the petrochemical route, the biotechnological production of lactate has several advantages, such as producing lactate with high enantiomeric purity, utilizing low-cost substrates, and promoting environmentally friendly processes (Alexandri et al., 2019). Industrially, about 90% of all lactate produced worldwide is derived from bacterial fermentation *via* pure homofermentative bacterial cultures in batch mode (Alves de Oliveira et al., 2018).

The use of open cultures of microbial consortia (*i*.*e*., reactor microbiomes) has economic and operational advantages, which includes the ability to use complex waste material as fermentation substrates without the need for sterilization (Policastro et al., 2021). Lactate production can be further improved by utilizing continuous processes, resulting in shorter fermentation times and higher production rates (López-Gómez et al., 2019). Additionally, continuous processes could prevent inhibition issues from high concentrations of substrate or their product (López-Gómez et al., 2019). The use of open-culture fermentation with reactor microbiomes has been unpopular, thus far, because of the challenges of preventing heterofermentation (Policastro et al., 2021), which can result in side products such as ethanol, acetate, and CO_2_ (Daly et al., 2020; Kandler, 1983). In addition, continuous lactate fermentation can experience problems related to unutilized sugars and cell wash-out, which are complications that are intensified when dilution rates are increased. Thus, cell recycling methods are necessary to achieve high cell densities to increase lactate productivity (López-Gómez et al., 2019). Previous studies have shown that a homofermentative microbiome that was dominated by *Lactobacillus* spp. can be enriched by maintaining a low pH and thermophilic conditions (Kim et al., 2016; Tang et al., 2017; Xu et al., 2018), and the lactate productivity can be enhanced when utilizing more effective cell recycling methods during continuous fermentations (Xu et al., 2018).

The dairy sector is the European Union’s (EU) second-biggest agricultural sector in economic output value (Walsh et al., 2022). Most of the milk in the EU is used for the production of cheese (37.7%) and butter (29.4%), and only 4.3% for acidified milk (Ghinea and Leahu, 2020). The milk conversion processes result in substantial amounts of wastewater that is rich in organic and non-organic compounds, posing considerable financial and technological challenges for the dairy industry (Walsh et al., 2022). Acid whey is one such wastewater from the dairy industry, which is generated during the production of acid-coagulated dairy products, such as fresh cheeses, cream cheese, or strained (Greek-style) yogurt, and from the production of calcium caseinate (Çelik and Yuksel, 2016; Kissalita et al., 1989). The EU has the world’s largest cheese market, and thus high acid-whey production, which is estimated to be around 40 million tonnes per year (Koller et al., 2005; Zotta et al., 2020). For 2019, the total acidified milk (yogurts and other) production in the European Union (EU) was estimated to be ∼8 million tonnes (Eurostat, 2020). During Greek yogurt production, ∼67% of the volume of milk is converted to acid whey (Rocha-Mendoza et al., 2021). While during acid-coagulated cheese production, ∼87% of the volume of milk is converted to acid whey (Carvalho et al., 2013).

Acid whey has a high biochemical oxygen demand (BOD) (30 to 60 g L^-1^) (Kissalita et al., 1989) and chemical oxygen demand (COD) (60 to 80 g L^-1^) (Cristiani-Urbina et al., 2000), categorizing it as high-organic wastewater that requires treatment before release into aquatic environments. The lactose and galactose within acid whey can be used as a carbon source for fermentation processes using lactic acid bacteria (LAB) to produce lactate as a sole product (Kissalita et al., 1989). Here, we investigated lactate production from acid-whey wastewater that was produced during Greek-style yogurt production. We enriched endogenous *Lactobacillus* spp., which originated from acid whey, in reactor microbiomes. We operated three reactor configurations with different methods of cell retention: **(1)** an upflow anaerobic sludge blanket (UASB) reactor; **(2)** an anaerobic filter reactor (AFR); and **(3)** an anaerobic continuously stirred tank reactor (CSTR) with a hollow-fiber membrane module to find the best reactor configuration for lactate production. We continuously fed acid whey to the three reactor microbiomes under acidic (pH 5.0 ± 0.1) and thermophilic conditions (50° ± 1°C). We tested different hydraulic retention times (HRTs) to find the operating parameters that would result in the highest lactate conversion rates and conversion efficiencies from the lactose- and galactose-rich acid-whey substrate.

## 2. Materials and methods

### 2.1. Acid whey

We utilized raw acid whey from Greek-style yogurt production as the substrate for the three reactor microbiomes. We collected acid whey at 13 different time points throughout the ∼1.5-year study from FrieslandCampina, GmbH (Cologne, Germany). We collected ∼300 L per batch in an IBC container (Richter & Heß Verpackungen, Chemnitz, Germany). Immediately upon arrival, we aliquoted the acid whey in 10-L plastic buckets and stored the aliquots at -20°C until further use. Before storage, we took liquid samples from three randomly chosen buckets to characterize the acid-whey batches (**Table S1**). The acid-whey aliquots were thawed in a warm water bath (40°-60°C) for 2-3 h before being used as the substrate for the three reactor microbiomes.

### 2.2. Bioreactor systems

The acid-whey substrate was stored in a glass-jacketed vessel and cooled at ∼2°-4°C using a recirculating water chiller (MiniChiller 600, Huber, Offenburg, Germany). We further insulated the substrate vessel with an AF/Armaflex 13-mm sheet (Armacell GmbH, Münster, Germany). We continuously mixed the acid-whey substrate by recirculating liquid from the top to the bottom of the bioreactor using a peristaltic pump that was equipped with Norprene^®^ Food I/P^®^ tubing (Masterflex, Cole-Parmer, St. Neots, UK). We used peristaltic pumps (Masterflex, Cole-Parmer, St. Neots, UK) that were equipped with BioPharm Plus^®^ I/P^®^ platinum-cured silicone tubing (Masterflex, Cole-Parmer, St. Neots, UK) to provide substrate to the reactor microbiomes continuously.

The upflow anaerobic sludge blanket (UASB) reactor consisted of a glass-jacketed column reactor with a height of 0.6 m, an inner diameter of 6 cm, and a wet volume of 2.8 L. We used a thermostatic water bath (CC-104A, Huber, Offenburg, Germany) to continuously recycle warm water through the jacket to heat the system to 50° ± 1°C. The reactor broth was continuously mixed in an upflow-hydraulic pattern at 4 m h^-1^ using a peristaltic pump that was equipped with Norprene^®^ Food I/P^®^ tubing. The influent was pumped into the recirculation line with a pump. We inserted a pH probe (SL80-225, SI Analytics GmbH, Mainz, Germany) through a dedicated port in the bioreactor lid. We connected the pH probe to an automated pH controller (Bluelab, Tauranga, New Zealand) that dosed a 2 M sodium hydroxide (NaOH) solution to regulate the pH at 5.0 ± 0.1. The effluent continuously exited the bioreactor through hydraulic forces *via* an overflow port at the top and was collected into an effluent reservoir.

We used an inverted glass funnel to collect the biogas that was produced within the bioreactor. The produced biogas was quantified with a gas flow meter (μFlow unit 2 mL resolution, BPC Instruments AB, Lund, Sweden). We included a biogas-collection system comprising a gas condenser, a sampling septum, a bubbler, and a gas-collection bag. On Day 308, we added 100 g of glass beads (Ø 10 ± 0.5 mm) (Carl Roth GmbH, Karlsruhe, Germany) and 40 g of pumice stone (granular, 1-5 mm) (Thermo Fisher Scientific, Waltham, MA) above the glass beads at the bottom of the UASB reactor for a better substrate distribution.

For the anaerobic filter reactor (AFR), we used a glass-jacketed column reactor with the exact dimensions of the UASB reactor. We filled the AFR with 322.2 g of Kaldnes K1 filter medium (Evolution Aqua, Wigan, UK), resulting in a wet volume of 2.2 L. The remaining setup of the AFR (pumps, mixing, temperature control, pH control, and biogas-collection system) did not differ from the UASB-reactor setup. We maintained an upflow hydraulic pattern with an upflow velocity of 2 m h^-1^ by recirculating mixed liquor with a peristaltic pump.

The anaerobic continuously stirred tank reactor (CSTR) consisted of a glass-jacketed reactor with a height of 0.2 m, an inner diameter of 11.4 cm, and a wet volume of 1.72 L. The reactor was closed with a custom stainless-steel lid. We installed a hollow-fiber membrane module (MiniKros Sampler 20 cm a 0.2 μm PES 1mm, Repligen GmbH, Ravensburg, Germany) to recycle the biomass back into the reactor. The permeate was pumped into an effluent reservoir. We recycled the reactor biomass at a flow of 160 mL min^-1^. We exchanged and cleaned the cell-recycle membrane weekly with a 2 M potassium hydroxide (KOH) solution. We equipped the CSTR with a stand-alone stirrer with a control box (Chemglass Inc, Vineland, NJ) and continuously stirred the reactor broth at 300 rpm. We connected a stainless-steel gas condenser to the biogas outlet. The biogas-collection system consisted of a sampling septum, an airless gas bubbler, a μFlow gas-flow meter, and a gas-collection bag. We connected the pH probe (SL80-120, SI Analytics GmbH, Mainz, Germany) through a dedicated port in the bioreactor lid. The remaining setup of the CSTR (pumps, tubing type, temperature control, and pH control) did not differ from the UASB-reactor and AFR setups.

### 2.3. Experimental periods

We did not use an external inoculum to start the bioreactors. Instead, we enriched the endogenous bacteria from the acid whey. We first operated the reactor in batch mode at a pH of 5.0 and a temperature of 44°C as part of the start-up phase. The start-up phase periods are described in the supplementary data, **S.1.1 Start-up phase periods**. Following the start-up phase, we tested the performance of the reactor microbiomes under the same operating conditions, which were a pH of 5.0 and a temperature of 50°C. We compared the abilities of the reactor microbiomes to convert acid whey into lactate under different hydraulic retention times (HRTs): **(1)** a 3-day HRT (Days 35-110, Days 29-104, and Days 28-103, for the UASB reactor, AFR, and CSTR, respectively); **(2)** a 2-day HRT (Days 110-184, 104-178, and 103-177, respectively); **(3)** a 1-day HRT (Days 184-273, 178-267, and 177-265, respectively); and **(4)** a 0.6-day HRT (Days 273-566, 267-560, and 265-559, respectively). We fed undiluted acid whey for the 3-, 2-, and 1-day HRTs and a 1.67-fold diluted acid whey for the 0.6-day HRT. The HRT periods discussed in this study and the performance parameters are described in **Table S2, Table S3**, and **Table S4**, for the UASB reactor, AFR, and CSTR, respectively.

### 2.4. Liquid sampling, analytical procedures, and calculations

We collected 1-mL liquid samples on a daily basis. We treated and stored the samples as described in Esquivel-Elizondo et al. (2021) until further analysis. We analyzed the samples using a high-performance liquid chromatography (HPLC) (LC20, Shimadzu Deutschland GmbH, Duisburg, Germany) system as previously described by Klask et al. (2020). We screened for the following compounds with the HPLC: **(1)** lactose (L); **(2)** galactose (G); **(3)** glucose (Glu); **(4)** lactate (LA); **(5)** ethanol (EtOH); **(6)** acetate (C2); and **(7)** *n*-butyrate (C4). We estimated the undissociated lactic acid (LA) concentrations *via* the Henderson–Hasselbalch equation (Harroff et al., 2017) as a function of measured pH values and the LA p*K*_*a*_ of 3.86. We tested the LA isomers proportions with a D-/L-Lactate (Rapid) Assay Kit (Megazyme, Bray, Ireland) (Schütterle et al., 2024). We analyzed the total (TS) and volatile (VS) solids according to Baird et al. (2017). We analyzed the total chemical oxygen demand (TCOD) and soluble chemical oxygen demand (SCOD) values with the closed reflux titrimetric method (Baird et al., 2017). We used a total organic carbon (TOC) analyzer (TOC-L Series, Shimadzu Deutschland GmbH, Duisburg, Germany) to analyze the TOC and dissolved organic carbon (DOC) as described by Schütterle et al. (2024). Before analyzing the COD and TOC, we homogenized samples with a T 10 basic ULTRA-TURRAX disperser (IKA-Werke GmbH & Co. KG, Staufen, Germany). Finally, we filtered the samples for SCOD and DOC through a 0.2-μm CHROMAFIL PET syringe filter (Carl Roth GmbH, Karlsruhe, Germany).

We calculated the reactor performance parameters in mmol C (Xu et al., 2018). The errors represent 95% confidence intervals throughout a steady-state period. It is important to note that when we mention the term conversion rate, we refer to the actual production rate of a specific product by excluding the influent concentration of that product (Xu et al., 2018). Moreover, we use the term production rate to refer to the total production, including the incoming influent concentration of that product. We report the equations that we used for calculating performance parameters in **S.2 Equations**. We performed a one-way ANOVA with the Tukey-Kramer test as the *post-hoc* analysis to compare the fermentation performance (*i*.*e*., volumetric LA conversion rate, LG-into-LA conversion efficiency, yield, and undissociated LA concentrations) between the reactor microbiomes at the different operating HRTs. We performed the statistical analyses in OriginPro, 2023 (OriginLab Corporation, Northampton, MA).

### 2.5. Biomass sampling, sequencing, and microbiome analysis

We collected biomass samples every week for microbial community characterization. Before sampling, we resuspended settled flocculent biomass in the mixed liquor of the UASB reactor and AFR by connecting a 60-mL syringe and quickly withdrawing and refilling the syringe ten times. We collected 2 mL of biomass sample in a sterile DNA- and Rnase-free 2.0-mL microcentrifuge tube (Carl Roth, Art. No. XC83.1) and centrifuged the samples for 6 min at 17,000 *g* in a benchtop centrifuge (5427 R, Eppendorf, Hamburg, Germany). We discarded the supernatant and stored the biomass pellets at -80°C until further analysis (as marked in **Fig. S1)**. Following the manufacturer’s protocol, we extracted the gDNA with the FastDNA™ SPIN Kit for Soil (MP Biomedicals, Irvine, CA) and assessed the resultant integrity *via* 1% agarose gel electrophoresis.

We targeted the V4 region of the 16S rRNA gene using universal primers 515F and 806R for the amplicon library construction. Libraries were constructed using dual index barcoding according to Schütterle et al. (2024). We sent the pooled library amplicons to the Genome Center at the Max Planck Institute (MPI) for Biology Tübingen, where the sequencing was carried out with the Illumina MiSeq Reagent Kit v2 (300-cycles). We deposited the raw .fastq files at the National Center for Biotechnology Information (NCBI) database under the accession number PRJNA884981.

We performed bioinformatic analysis with QIIME2 v2022.2 (Bolyen et al., 2019). Raw sequence data were demultiplexed and quality filtered using the q2-demux plugin followed by denoising with DADA2 (Callahan et al., 2016), *de-novo* clustering at 99% identity (Rognes et al., 2016), and removal of operational taxonomic units (OTUs) found in less than three samples. We assigned the taxonomy using the classify-sklearn naïve Bayes taxonomy classifier plugin (Bokulich et al., 2018) against the SILVA 138.1 database (Quast et al., 2013) using a RESCRIPt classifier for the V4 region (Robeson et al., 2021). We removed sequences not classified at the kingdom level or when classified as Eukaryota, Archaea, chloroplasts, or mitochondria. We used mafft alignment (Katoh et al., 2002) and fasttree2 (Price et al., 2010) to construct a rooted tree that we used for subsequent phylogenetic diversity analyses. Samples were rarefied at a sampling depth of 35,804 sequences per sample. We analyzed the α diversity *via* the Shannon diversity index, observed OTUs, and dominance metrics for the different operating HRTs. We performed Kruskal-Wallis tests for pairwise comparison, followed by a *post-hoc* analysis with Dunn’s test and Benjamini-Hochberg (BH) FDR adjustment if there were significant differences (*p* < 0.05).

The weighted and unweighted UniFrac distance metrics were used to analyze β diversity (Lozupone and Knight, 2005; Lozupone Catherine et al., 2007). We carried out a distance-based redundancy analysis (dbRDA) using the Bray–Curtis dissimilarity with the capscale function from the vegan package with a square root transformation and Wisconsin double-standardization (Oksanen et al., 2016) in RStudio v2022.12.0.353 (RStudio Team, 2022). To visualize the ordination, we correlated the most significant genera (>1% relative abundance) with the assemblage variation and displayed them as eigenvectors to indicate the strength and direction of the relationships (Lindfield et al., 2016). The envit function was used to obtain the *p*-value of correlation for the vector selection. We used the ggplot2 package (Wickham, 2021) to visualize the outcome. Finally, we created heat maps to represent the OTU relative abundance at the genus level using OriginPro, 2023 (OriginLab Corporation, Northampton, MA).

## 3. Results and discussion

### 3.1. Incomplete conversion of lactose and galactose to lactate was observed among the three reactor configurations

Xu et al. (2018) observed higher conversion efficiency of the acid-whey sugars, lactose and galactose, to lactate after changing their cell retention method. Xu et al. (2018) replaced the gravity settler in their CSTR with a hollow-fiber membrane, and they observed a 1.7-fold increase in the conversion efficiency due to the higher biomass retention efficiency. In this study, we operated the three reactor configurations (*i*.*e*., UASB reactor, AFR, and anaerobic CSTR with a hollow-fiber membrane module) with different biomass retention methods to ascertain the most promising reactor configuration for acid whey to lactate conversion. Similarly to Xu et al. (2018), during the comparison periods to achieve lactate homofermentation, we operated the three reactors at a pH of 5.0 and a temperature of 50°C. Contrary to Xu et al. (2018), we did not use an external inoculum to start the reactors. Instead, we enriched the lactate-producing microbiome from endogenous bacteria within the acid whey during the start-up phase periods (see supplementary data, **S.1.1 Start-up phase periods**)

The acid whey that we used in this study had similar properties (**Table S1**) to the acid whey that Xu et al. (2018) used in their study, with the main components being lactose, galactose, and lactate (*i*.*e*., calculated SCOD* using the main components concentrations from HPLC analysis-to-experimental SCOD ratio of 0.90 ± 0.03). The acid-whey batches that we collected and used in this study were very similar with TCOD and SCOD concentrations of 69.6 ± 1.62 and 65.7 ± 1.60 g COD L^-1^. We observed that the acid-whey composition in the reactor substrate tank and lines could change and differ from the acid whey upon arrival, with ethanol and acetate being formed; therefore, we minimized this process and regularly sampled the acid-whey substrate tank to have the correct volumetric organic loading rate composition (**Table S2, Table S3**, and **Table S4**). In addition to studying the effect of reactor configurations on the lactate conversion rates and LG-into-LA conversion efficiencies, we further tested the impact of the HRT and the subsequent substrate loading change by reducing the HRT from 3 days to 0.6 days (**Fig. 1**). Indeed, we observed significant differences (*p* < 0.05, Tukey-Kramer, **Fig. 2A-P**) in performance parameters among the three reactors during the different HRT periods, with exception to the lactate yield at the 3-day HRT where there was no significant difference (**Fig. 2I**, *p* **=** 0.956, Tukey-Kramer).

**Fig 1.**
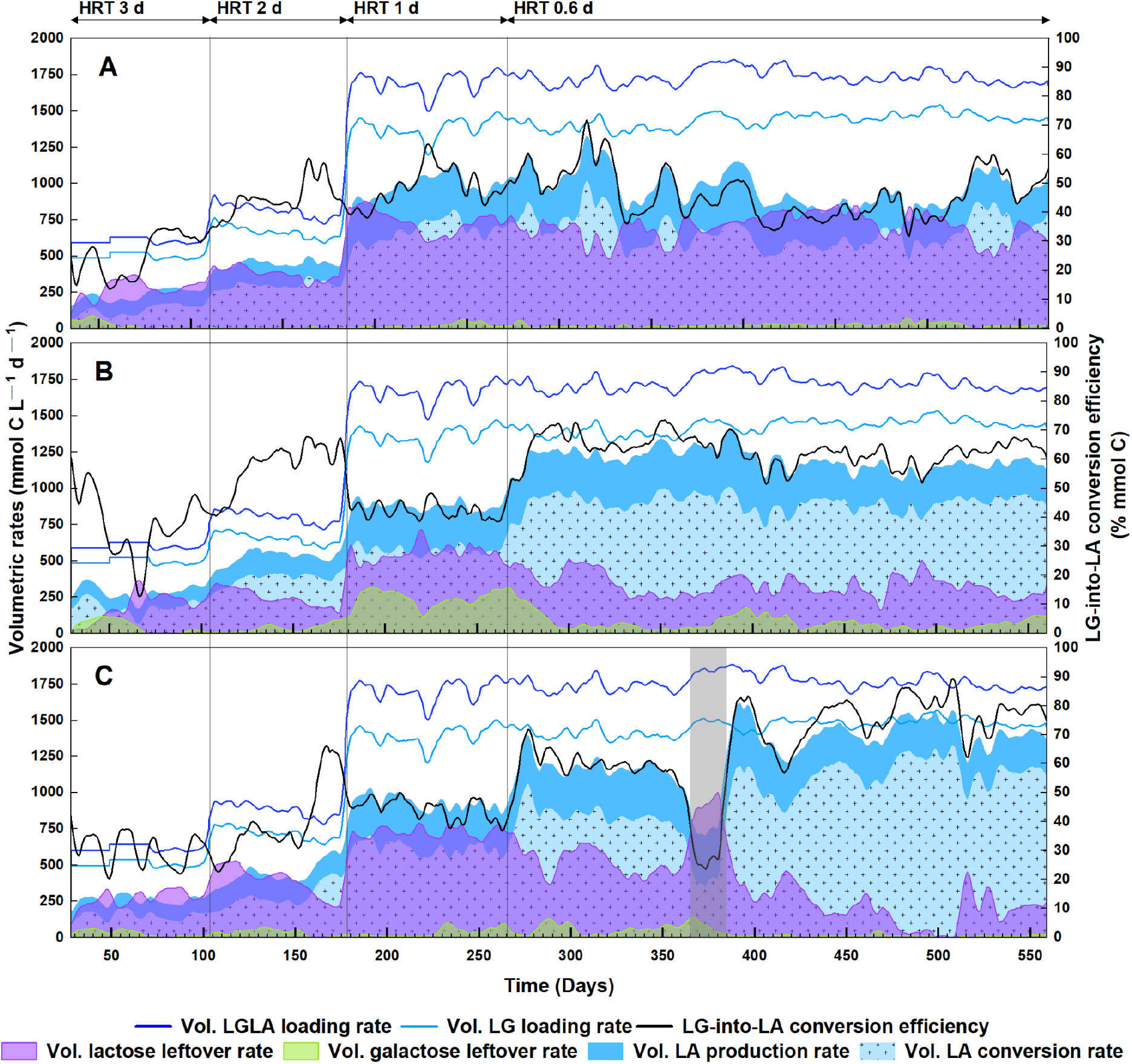
Performance of the reactor microbiomes during the different operating HRTs. (A) UASB reactor; (B) AFR; and (C) CSTR. Area plots of the lactate production rate, lactate conversion rate, and leftover substrate rates (lactose and galactose) on the left axes. Line plots of the total organic loading rate (lactose, galactose, and lactate) and the substrate loading rate (lactose and galactose) on the left axes. Line plots of the LG-into-LA conversion efficiencies are shown on the right axes. The data represent a 6-day moving average (each average contains six sampling points). The time period for the CSTR marked with gray represents the pH-error period when the actual pH value was 4.3. Vol. in volumetric.

**Fig 1.**
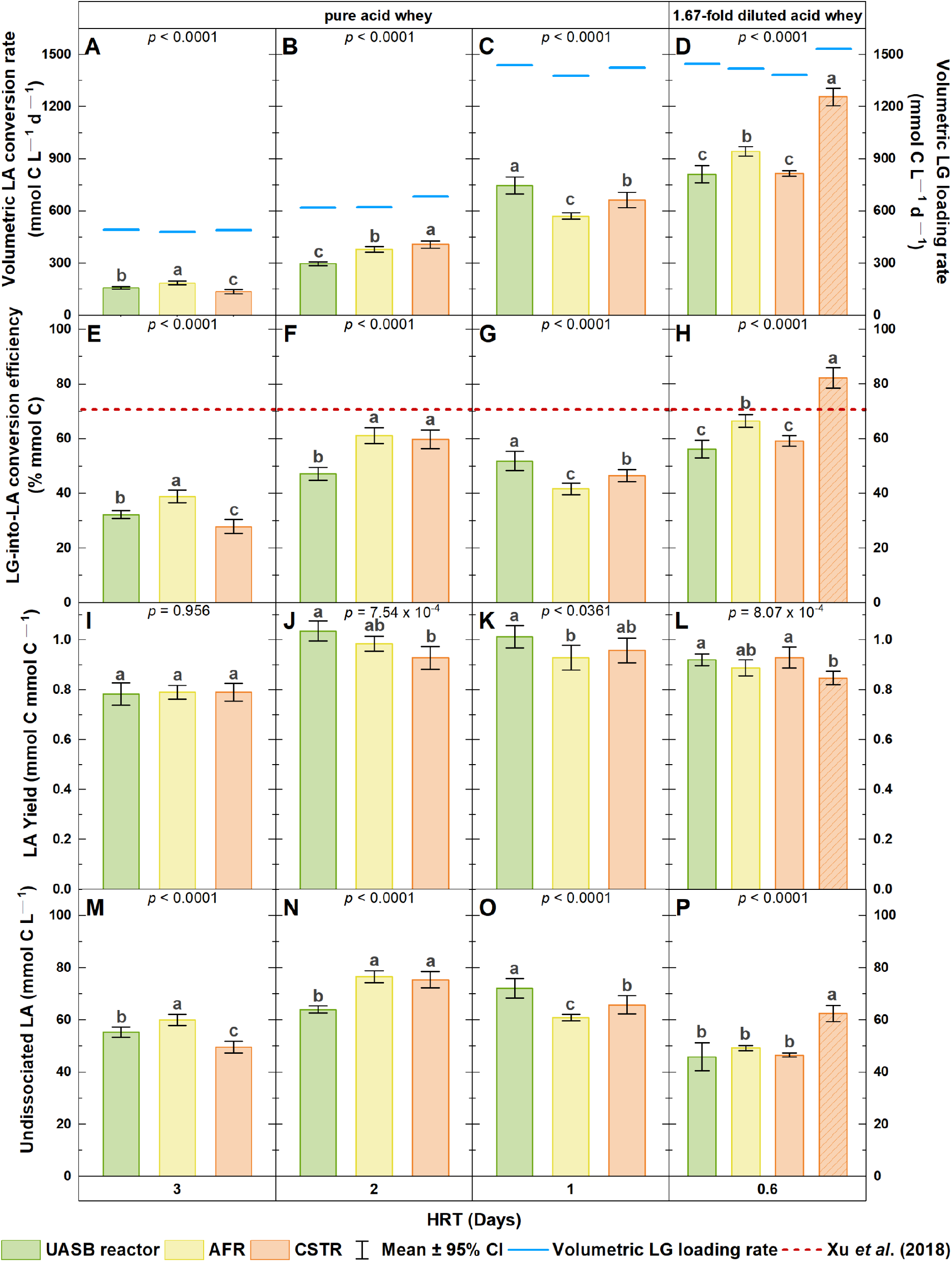
Comparisons of the reactor microbiomes’ performance parameters during the different operating HRTs. (A-D) lactate conversion rate; (E-H) substrate (lactose and galactose) to lactate conversion efficiency; (I-L) lactate yield based on substrate consumption; and (M-P) undissociated lactic acid concentration. Bar plots represent means between ≥ 15 days of steady-state operation conditions at a given HRT. At the 0.6-day HRT for the CSTR, the bar without a pattern represents data before the pH-error period, and the patterned bar represents data after the pH-error period. One-way ANOVA with Tukey-Kramer′s test as the *post-hoc* analysis was used to test significant differences. Significant differences (*p* < 0.05) between the reactor microbiomes at a given HRT are indicated with different letters. For means denoted with the same letter, the difference between the means is not statistically significant (*e*.*g*., in panel J, the significance letters indicate that there is no significant difference between the UASB reactor and the AFR and between the AFR and the CSTR. However, there is a significant difference between the UASB reactor and the CSTR). (A-D) Scattera plots of the volumetric LG loading rate on the right axes. (E-H) The dashed red line shows the conversion efficiency obtained by Xu et al. (2018).

**Fig 2.**
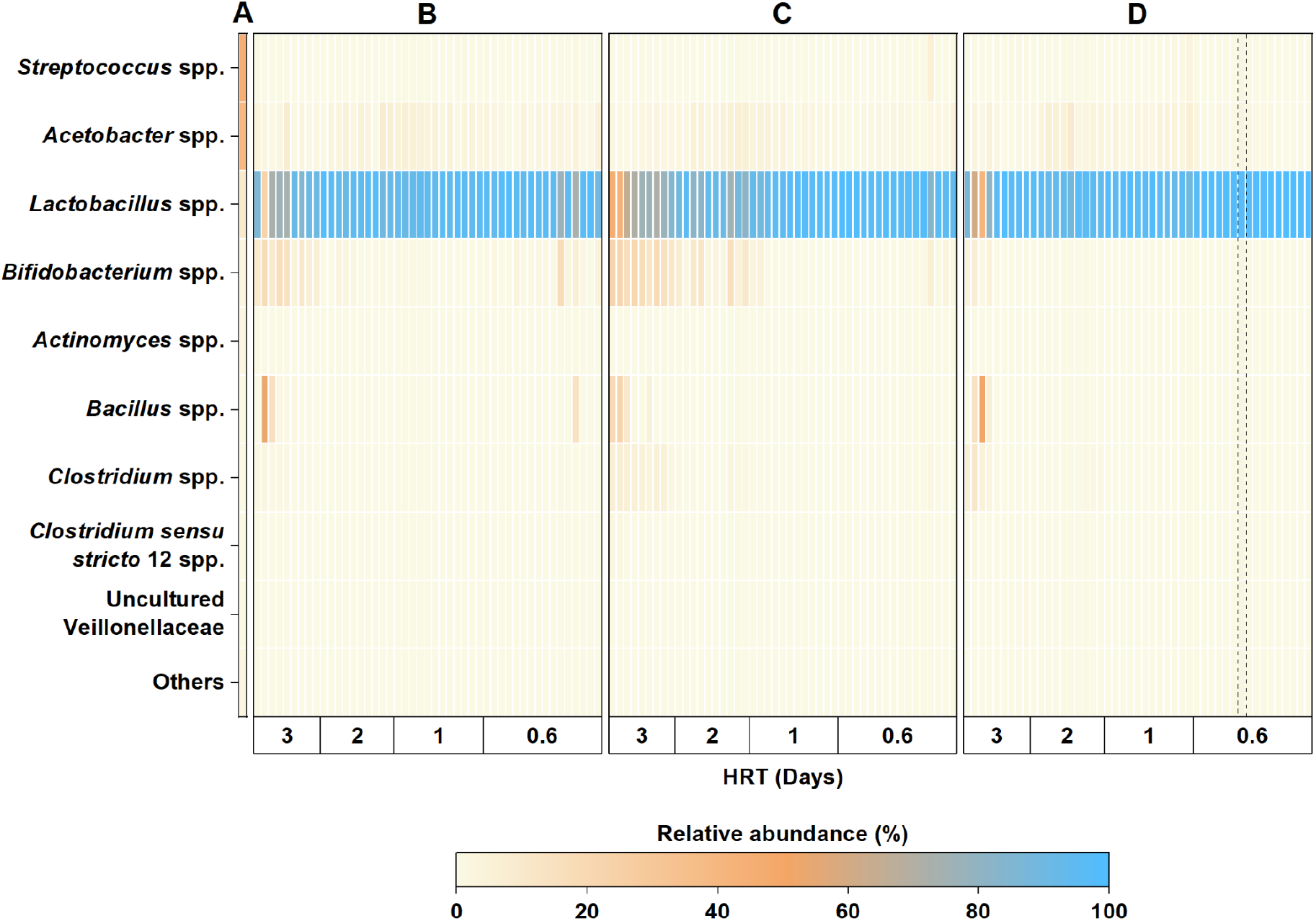
Heat map of relative genera abundances at Time 0 and during the different operating HRTs. (A) the acid-whey batch that was used to autoinoculate the bioreactors; (B) UASB reactor; (C) AFR; and (D) CSTR. Genera that were abundant < 1% across different samples were summed in Others. The marked period at the 0.6-day HRT for the CSTR represents samples taken when the pH control malfunctioned, and the actual pH value was 4.3.

During the 3-day HRT, the AFR had a significantly higher lactate conversion rate and LG-into-LA conversion efficiency (**Fig. 2A, E**, *p* < 0.0001, Tukey-Kramer) than the UASB reactor and CSTR. The lactate conversion rates were 158 ± 6.72, 186 ± 10.4, and 136 ± 11.0 mmol C L^-1^ d^-1^ for the UASB reactor, AFR, and CSTR, respectively (**Fig. 2A**). Nevertheless, the LG-into-LA conversion efficiencies (**Fig. 2E**) were insufficient in all three reactors, with values < 40% (in mmol C), resulting in high amounts of leftover substrate (**Fig. 1**). The LG-into-LA conversion efficiencies were 32.8 ± 1.38, 38.9 ± 2.23, and 27.9 ± 2.38% for the UASB reactor, AFR, and CSTR, respectively (**Fig. 2E**). These results were far below our goal to surpass the LG-into-LA conversion efficiency of 70.7% that Xu et al. (2018) had achieved. In all three reactors, we observed the production of the side products (**Fig. S1**), ethanol, acetate, and *n*-butyrate, with side product yields of 0.04 ± 0.01, 0.05 ± 0.01, and 0.05 ± 0.02 mmol C mmol C^-1^ for the UASB reactor, AFR, and CSTR, respectively (**Table S2, Table S3**, and **Table S4**). The lactate yields were 0.78 ± 0.04, 0.79 ± 0.03, and 0.79 ± 0.03 mmol C mmol C^-1^, respectively (**Fig. 2I**). The undissociated lactic acid concentrations within the reactors were 55.3 ± 1.94, 60.0 ± 2.00, and 49.6 ± 2.20 mmol C L^-1^, respectively (**Fig. 2M**). The low LG-into-LA conversion efficiencies and incomplete substrate utilization indicated that product inhibition was occurring in the reactors, specifically due to the undissociated acid form (Honda et al., 1995; Iyer and Lee, 1999; Venkatesh et al., 1993; Yabannavar and Wang, 1987).

Shortening the HRT to 2 days resulted in: **(1)** higher lactate conversion rates (297 ± 9.87, 380 ± 14.1, and 408 ± 19.7 mmol C L-1 d^-1^, respectively) (**Fig. 2B)**; **(2)** higher LG-into-LA conversion efficiencies (47.2 ± 2.16, 61.2 ± 2.64, and 59.8 ± 3.17%, respectively) (**Fig. 2F**); and **(3)** lower side product yields (< 0.01, 0.03 ± 0.01, and 0.02 ± 0.00 mmol C mmol C^-1^, respectively) (**Table S2, Table S3**, and **Table S4**), for the UASB reactor, AFR, and CSTR, respectively. At this HRT, the lactate yields increased to 1.04 ± 0.04, 0.98 ± 0.03, and 0.93 ± 0.04 mmol C mmol C^-1^, respectively (**Fig. 2J**), while the undissociated lactic acid concentrations in the reactors increased to 64.3 ± 1.34, 76.5 ± 2.03, and 75.5 ± 2.85 mmol C L^-1^, respectively (**Fig. 2N**). The faster product wash-out probably helped to partially alleviate the inhibiting effect and shift the inhibition limit for the undissociated acid form, which could explain the increase of the LG-into-LA conversion efficiencies.

When we further reduced the HRT to 1 day, we observed a drop in the LG-into-LA conversion efficiency for the AFR and CSTR (41.7 ± 1.91% and 46.5 ± 2.03%, respectively) (**Fig. 2G**), while for the UASB reactor, the LG-into-LA conversion efficiency (51.9 ± 3.17%) (**Fig. 2G**) remained similar compared to the 2-day HRT (**Fig. 2F**). The lactate yields were 1.01 ± 0.04, 0.93 ± 0.05, and 0.96 ± 0.05 mmol C mmol C^-1^ (**Fig. 2K**), while the side product yields were < 0.01, 0.02 ± 0.00, and < 0.01 mmol C mmol C^-1^ (**Table S2, Table S3**, and **Table S4**), for the UASB reactor, AFR, and CSTR, respectively. During this HRT, the undissociated lactic acid concentrations in the reactors reached 72.2 ± 3.45, 60.9 ± 1.11, and 65.8 ± 3.21 mmol C L^-1^, respectively (**Fig. 2O**). At the 2-day HRT and 1-day HRT, the increase of the lactate yield above 0.90 mmol C mmol C^-1^, indicates that lactate was produced due to non-growth (maintenance) associated production (Venkatesh et al., 1993), which could explain the subsequent drop of the conversion efficiencies at the 1-day HRT, due to the higher cell wash-out rate.

### 3.2. Undissociated lactic acid concentrations limited the acid-whey conversion potential of the reactor microbiomes

At lower pH values of 5.0 in our reactors, lactate production was inhibited by the undissociated acid form of lactate (Aghababaie et al., 2015) and most of the lactate was formed due to a non-growth (maintenance) associated production (Venkatesh et al., 1993). We speculated that undissociated lactic acid concentrations in the range ∼50-60 mmol C L^-1^ (**Fig. 2M**) were inhibiting the reactor microbiomes because even at the longest 3-day HRT that we operated (**Fig. 1, Table S2, Table S3**, and **Table S4**), we observed less than 50% of carbon utilization based on mmol C. To test whether the undissociated lactic acid concentration was causing incomplete substrate conversion, we further decreased the HRT from 1 day to 0.6 days. However, we diluted the acid whey 1.67-fold to achieve the same loading rate at the 0.6-day HRT (**Fig. 2D**) compared to the 1-day HRT (**Fig. 2C**). We hypothesized that the LG-into-LA conversion efficiencies would increase if we maintained the undissociated lactic acid concentration below ∼ 60 mmol C L^-1^.

Indeed, we observed an increase in the LG-into-LA conversion efficiencies compared to the 1-day HRT. We achieved lactate conversion rates of 812 ± 45.1 and 942 ± 24.6 mmol C L^-1^ d^-1^ and LG-into-LA conversion efficiencies of 56.1 ± 2.98% and 66.5 ± 2.17% for the UASB reactor and AFR, respectively (**Fig. 2D, H**). In addition, we achieved a lactate conversion rate of 817 ± 19.8 mmol C L^-1^ d^-1^ and an LG-into-LA conversion efficiency of 59.2 ± 1.80% for the CSTR before Day 365 (**Fig. 1**). During the period between Days 365 and Day 384 (**Fig. 1**), we experienced a fault in the pH control of the CSTR system. Due to a faulty pH probe, the pH in the CSTR dropped to a pH value of 4.3 during this period. The undissociated lactic acid concentration reached 118 ± 7.83 mmol C L^-1^. After we replaced the pH probe, the CSTR immediately recovered. The lactate conversion rate and LG-into-LA conversion efficiency increased to 1256 ± 46.3 mmol C L^-1^ d^-1^ (1.57 ± 0.06 g L^-1^ h^-1^) and 82.2 ± 3.41%, respectively (**Fig. 2D, H**), surpassing the UASB reactor and AFR and the results obtained by Xu et al. (2018). The LA-to-SCOD effluent concentration ratio in the CSTR reached 96.9% g COD (**Table S4**). The exposure to high undissociated lactic acid concentrations for an extended period could have allowed the microbiome to adapt and achieve higher tolerance. In the study where whey was utilized as a substrate for continuous reactor microbiomes, Choi et al. (2016) observed an improvement in the lactate yield after the pH was increased to 5.5 in a reactor that was subjected to a pH value of 3.0 at the beginning of the fermentation.

Using diluted acid whey as substrate (**Fig. 2D**) kept the undissociated lactic acid concentration (**Fig. 2P**) in the reactors below the inhibition level, and operating at shorter HRTs allowed for high lactate conversion rates (**Fig. 2H**). Due to the non-growth associated lactate production, the conversion rates were dependent on the biomass concentration and the ability of the reactor configuration to retain biomass. We did not observe the formation of typical granulated biomass in the UASB reactor throughout the entire operating period. Only on Day 380 did we observe a fluffy floc formation, which is typical for wastewaters with a relatively high protein content (Thaveesri et al., 1994). These fluffy flocs can be easily lost (Thaveesri et al., 1994), which we observed in the UASB reactor after a week. The inability to form granular biomass and efficiently retain cells could explain the underperformance of the UASB reactor at the high dilution rates of 1.67 d^-1^ compared to the AFR and CSTR, where biomass was more efficiently retained. In this study, we used NaOH to control the pH to 5.0. The consumption of NaOH (**Eq. S1**) in all reactors was close to the theoretical value of 0.44 gNaOH_used_ g^-1^ LA_converted_ (**Table S2, Table S3**, and **Table S4**), which was also observed in previous studies (Acuña et al., 1994; Zheng et al., 2017).

### 3.3. *The acidic and thermophilic conditions enriched a specialized homofermentative microbiome dominated by* Lactobacillus *spp*

The most abundant population in the acid-whey substrate, which encompassed the microbes that autoinoculated the reactors, was *Streptococcus* spp. with a relative abundance of 45.1% (**Fig. 3A**). The second most abundant population in the acid whey was *Acetobacter* spp. with 38.7% relative abundance. *Lactobacillus* spp. was present at a 9.3% relative abundance in the acid whey. This changed drastically throughout the operating periods in the reactor microbiomes where *Lactobacillus* spp. was the most dominant genus (**Fig. 3B, C**, and **D**). During the 3-day HRT, *Lactobacillus* spp. dominated the microbiome at 76.5 ± 19.9%, 67.7 ± 13.5%, and 83.9 ± 18.9% relative abundance for the UASB reactor, AFR, and CSTR, respectively (**Fig. 3B, C**, and **D**). The second most abundant genus in the UASB reactor and AFR was *Bifidobacterium* spp. at a 10.7 ± 4.58% and 17.3 ± 4.07% relative abundance. For the CSTR, *Bifidobacterium* spp. were only present at a 1.95 ± 2.52% relative abundance during the 3-day HRT. The second most abundant genus for the CSTR at the 3-day HRT was *Bacillus* spp. at a 7.87 ± 15.7% relative abundance.

**Fig 3.**
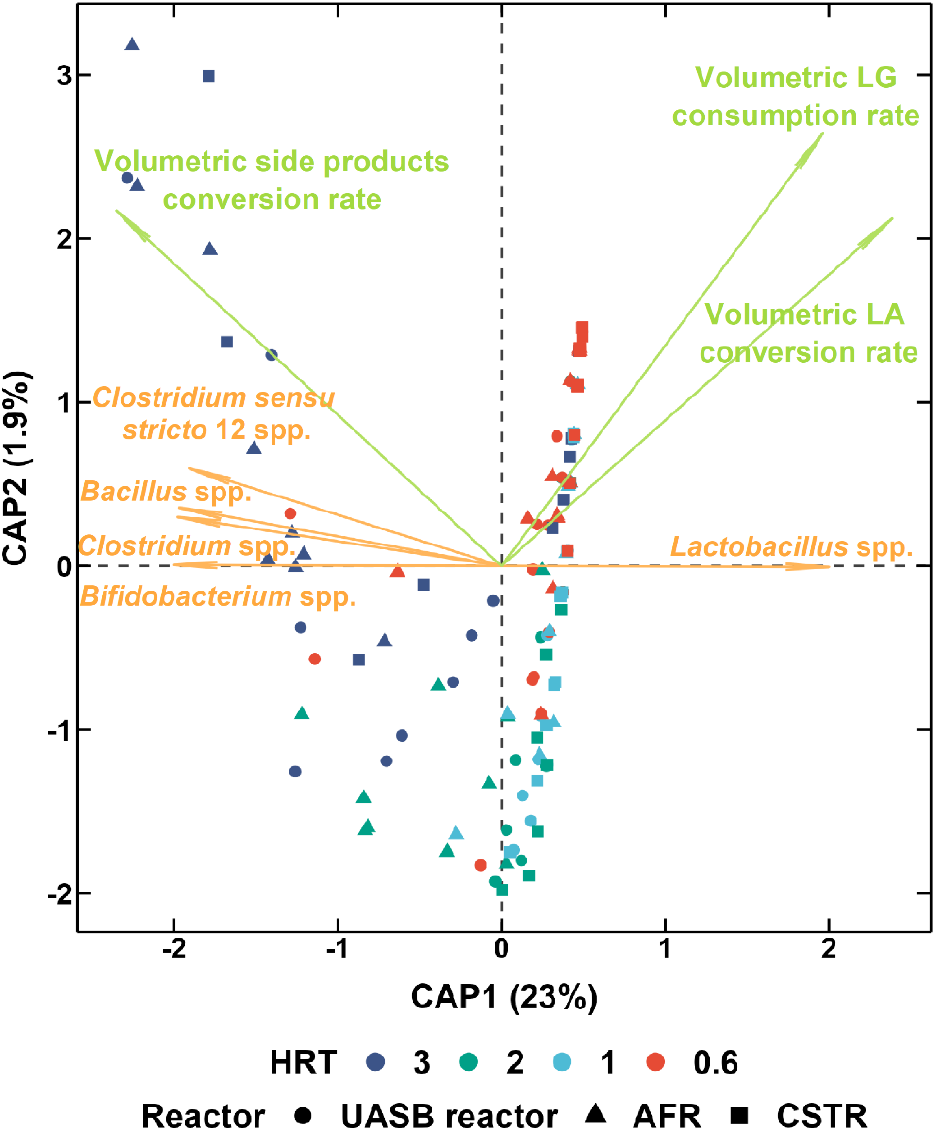
Distance-based redundancy analysis (dbRDA) of β diversity with Bray-Curtis squared dissimilarity. Vector ordinations show the strength and direction of the relationships for genera (orange) and derived kinetic process parameters (light green).

We observed that with the shortening of the HRT, *Lactobacillus* spp. became more dominant (**Fig. 3B-D**, and **Fig. S2G-I**). At the final HRT of 0.6 days, *Lactobacillus* spp. dominated the reactor microbiomes of the UASB reactor, AFR, and CSTR at 93.0 ± 7.05%, 96.1 ± 3.53%, and 98.6 ± 0.77% relative abundance, respectively. Before the pH error on Day 365-384 (**Fig. 1**) in the CSTR at the 0.6 day-HRT, *Lactobacillus* spp. was dominant at a 98.2 ± 0.65% relative abundance. After the pH-error period, the relative abundance was similar at 98.9 ± 0.72%. More specifically, *Lactobacillus* spp. comprised of two OTUs with 100% identity similarity to *Lactobacillus delbrueckii* subsp. *lactis* and *Lactobacillus acidophilus* in the NCBI database based on their 16S rRNA genes. More specifically, before the pH-error period, *L. delbrueckii* subsp. *lactis* and *L. acidophilus* were present in the CSTR at a 97.2 ± 0.90% and 1.01 ± 0.34% relative abundance, respectively. These values remained similar after the pH-error period, with *L. delbrueckii* subsp. *lactis* and *L. acidophilus* being present at a 97.7 ± 1.09% and 1.28 ± 0.48% relative abundance, respectively. Because of these very high abundances, we postulate that the pH-error period resulted in adaptive evolution of the strains, making them more tolerant to acid stress, and thus increasing the LG-into-LA conversion efficiency. Zhang et al. (2012) improved the acid tolerance in *L. casei* by adaptive evolution in their study with a duration of 70 days. The evolved *L. casei* mutant exhibited higher intracellular pH, NH4^+^ concentration, lower inner membrane permeability, and higher amounts of intracellular aspartate and arginine, resulting in a higher tolerance to acid stress. The biomass recirculation in our CSTR could have had a positive effect and reduced the necessary time for evolution to be achieved.

There were significant differences (*p* < 0.05, Kruskal-Wallis Test) in the α-diversity metrics (*i*.*e*., **(1)** Shannon index; **(2)** observed OTUs; and **(3)** dominance) for the individual reactor microbiomes between the different operating periods (**Fig. S2**), and between the reactor microbiomes at HRTs of 3, 2, and 0.6 days (**Fig. S3**). There were no significant differences for the 1-day HRT between the different reactor microbiomes for the Shannon index (**Fig. S3C**) and dominance (**Fig. S3K**) metrics. At the 3-day HRT, the Shannon index was 1.53 ± 0.55, 2.18 ± 0.40, and 1.46 ± 0.38 for the UASB reactor, AFR, and CSTR, respectively (**Fig. S3A**). The number of observed OTUs was 16.7 ± 4.03, 24.3 ± 4.75, and 15.6 ± 4.00, respectively (**Fig. S3E**). We observed a loss in diversity with a decrease in the Shannon index and the number of observed OTUs, and with an increase in the dominance metric after we shortened the HRT from 3 days to 0.6 days. At the 0.6-day HRT, the Shannon index was 0.55 ± 0.37, 0.50 ± 0.26, and (0.25 ± 0.07 and 0.21 ± 0.09) for the UASB reactor, AFR, and CSTR (before and after the pH-error period), respectively (**Fig. S3D**). At the 0.6-day HRT, the number of observed OTUs were 14.9 ± 1.61, 15.1 ± 2.84, and (11.7 ± 1.21 and 10.4 ± 2.46), respectively (**Fig. S3H**). The dominance metrics increased from 0.55 ± 0.18, 0.34 ± 0.11, and 0.50 ± 0.11, respectively at the 3-day HRT (**Fig. S3I**) to 0.86 ± 0.12, 0.88 ± 0.07, and (0.94 ± 0.02 and 0.95 ± 0.02), respectively at the 0.6-day HRT (**Fig. S3L**) for the UASB reactor, AFR, and CSTR (before and after the pH-error period at the 0.6-day HRT). All three microbiomes at the 0.6-day HRT were less diverse than the reactor microbiome operated by Xu et al. (2018) at a 1-day HRT in a CSTR, where the Shannon index was 0.55 ± 0.27, and the number of observed OTUs was 78 ± 4.

We performed a PERMANOVA analysis and observed significant differences in the β-diversity weighted UniFrac distance metrics between the reactor microbiomes in the UASB reactor, AFR, and CSTR at all HRTs (*p* = 0.001 at all HRTs, except for the 1-day HRT where *p* = 0.039). The pairwise PERMANOVA test at the 1-day HRT showed that there was a significant difference (*p* = 0.014) among the UASB reactor and CSTR population at the 1-day HRT, but not between the UASB reactor and AFR (*p* = 0.204), and between the AFR and CSTR (*p* = 0.122). At the 0.6-day HRT, we observed no difference in the population composition between the UASB reactor and AFR (*p* = 0.072). However, the CSTR population differed significantly (*p* = 0.001) from the UASB reactor and AFR at the 0.6-day HRT. There was a significant difference (*p* = 0.045) in the CSTR microbiome population before and after the pH-error period based on the weighted UniFrac. A different reactor microbiome is another explanation for the adaptation of the microbiome, besides strain evolution as mentioned above, to explain why the CSTR outperformed the UASB reactor and the AFR. Our resolution with 16S rRNA gene sequencing is not sufficient to explore the true reason for this adaptation during the pH error period.

The dbRDA analysis (**Fig. 4**) of all reactor microbiome samples at the different operating HRTs showed a positive correlation of the relative abundance of the homofermentative *Lactobacillus* spp. with the conversion of lactose and galactose to lactate. Furthermore, the dbRDA analysis correlated species of the genera *Bifidobacterium, Bacillus, Bifidobacterium, Clostridium*, and *Clostridium sensu stricto* 12 with side product formation. We observed from the dbRDA analysis that the shortening of the HRT led to an increase in the dominance of *Lactobacillus* spp.. We hypothesize that: **(1)** the ability of *Lactobacillus* spp. to thrive in acidic environments (Bendig et al., 2023; Schütterle et al., 2024); **(2)** their thermotolerance (Tashiro et al., 2011; Walters et al., 2024); and **(3)** the ability to produce bacteriocins that have antimicrobial properties against other Gram-positive bacteria or closely related microbes (Mora et al., 2020), gave them an advantage over other microbes, which resulted in the enriched *Lactobacillus* spp. microbiomes.

**Fig 4.**
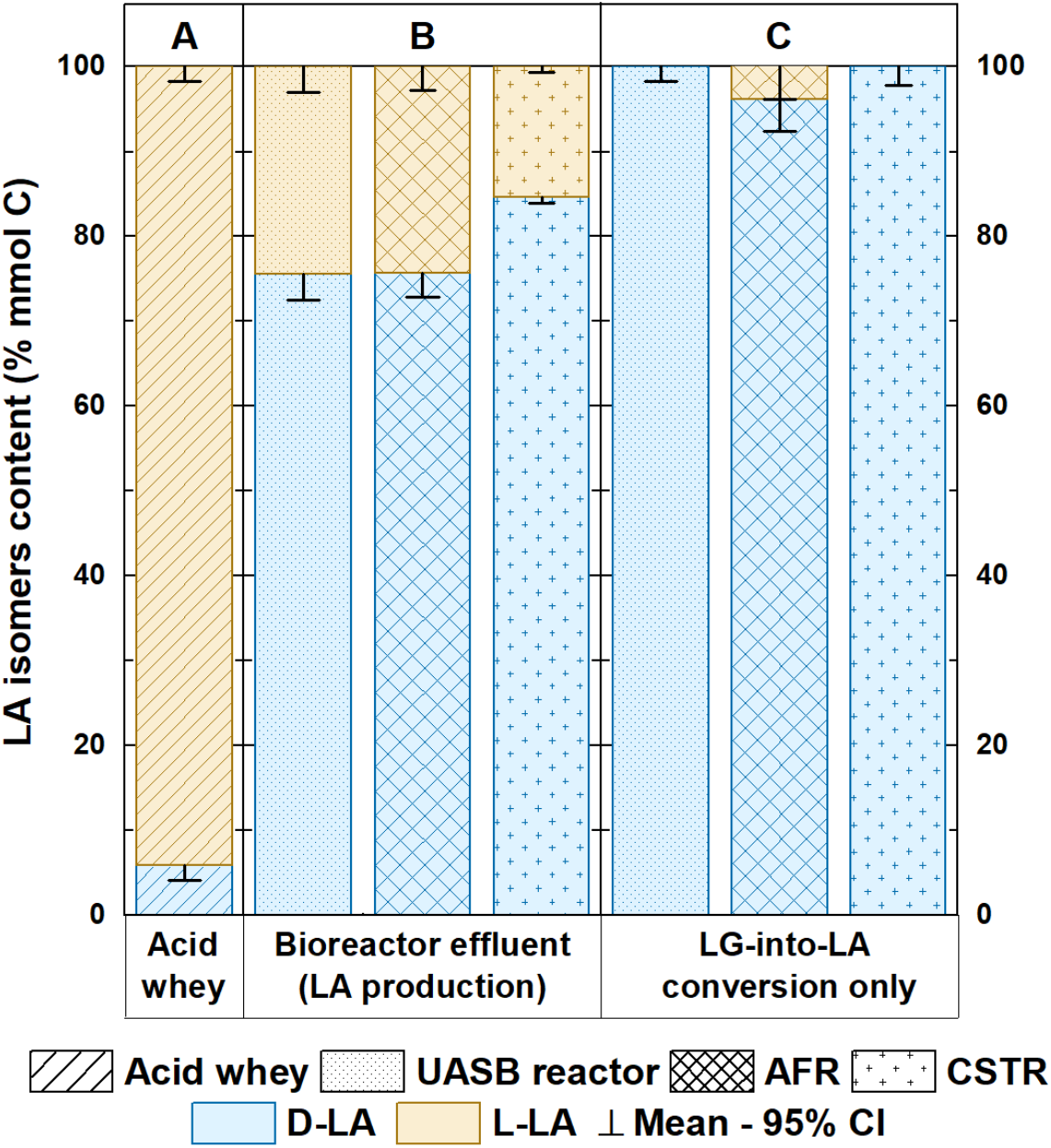
The lactate isomer proportions change after fermentation at the 0.6-day HRT. Bar plots of mean values (*n* = 3) of the lactate isomers present in the (A) acid whey; (B) bioreactor effluent; and (C) lactate conversion rate only for the UASB reactor (Days 541, 542, and 548), AFR (Days 546, 549, and 554), and CSTR (Days 548, 550, and 556). All sampling points were tested in triplicates.

To achieve higher conversion rates at shorter HRTs, we recommend an AFR or a CSTR with a hollow-fiber membrane module to prevent cell wash-out rather than a UASB reactor, which did not grow a sustainable granular bed. As mentioned earlier, retaining biomass is paramount for homofermentation of lactate due to non-growth (maintenance) associated production. Before the pH-error period for the CSTR, the AFR with a more straightforward internal biomass retention system outperformed the CSTR (**Fig. 1B-C**). Unfortunately, only an unintended and extensive period of low pH occurred for the CSTR. Not for the AFR (**Fig. 1**), which led to a change in the reactor microbiome composition in the CSTR only (**Fig. 4**), possibly in tandem with strain evolution, resulting in a considerable improvement in its performance (**Fig. 1C**). As a result, the CSTR with cell retention module was the superior bioreactor configuration. However, we predict that when the AFR would have gone through a similar low pH period, adapting the reactor microbiome, the AFR would have outperformed the CSTR. This is speculation, but if correct, the AFR would have been the preferred bioreactor configuration due to the less complex cell retention mechanism of having a matrix inside the bioreactor rather than an external cell retention module for the CSTR. Regardless, we recommend a low pH period as part of the start-up strategy to adapt the reactor microbiome.

### 3.4. Production of D-lactate from acid whey via reactor microbiomes

We investigated which lactate isomer from the reactor microbiomes was produced at the 0.6-day HRT. We observed that the homofermentative microbiomes converted lactose and galactose to exclusively D-lactate (**Fig. 5C**). The D-lactate isomer was represented with 100 ± 1.75%, 96.2 ± 3.77%, and 100 ± 2.13% as part of the LG-into-LA conversion rate for the UASB reactor, AFR, and CSTR, respectively (**Fig. 5C**). The presence of the L-lactate in the bioreactor effluent (*i*.*e*., lactate production rate) was 24.4 ± 3.04%, 24.4 ± 2.77%, and 15.4 ± 0.67% for the UASB reactor, AFR, and CSTR, respectively (**Fig. 5B**). This was due to L-lactate isomer that was already present in the acid whey substrate (**Fig. 5A**). The L-lactate isomer was the primary form present in the acid whey (94.1%) (**Fig. 5A**), which could be explained by the dominance of *Streptococcus* spp. in the acid whey (**Fig. 3A**). *Streptococcus* spp. are known L-lactate producers (Panesar et al., 2007). Commonly used microbes in yogurt production are the L-lactate producer *Streptococcus thermophilus* and the D-lactate producer *Lactobacillus bulgaricus* (Panesar et al., 2007). The operating parameters in our study allowed us to enrich the D-lactate producers used in yogurt production. Our work confirmed that a pH of 5.0 and a temperature of 47°-50°C were necessary to produce exclusively D-lactate from acid whey with reactor microbiomes (Schütterle et al., 2024). Our results were consistent with previous literature and showed that operating reactor microbiomes at thermophilic conditions creates a specialized reactor microbiome to perform lactate homofermentation (Kim et al., 2016; Xu et al., 2018). This homofermentative-specialized microbiome is characterized with non-growth (maintenance) associated production, which makes the need for cell retention a priority. To achieve higher conversion rates at shorter HRTs, we recommend an AFR or an anaerobic CSTR with a hollow-fiber membrane module, or another reactor configuration that prevents cell wash-out.

## 4. Conclusion

- A pH of 5.0 and temperature of 50°C enriched the *Lactobacillus* spp. that were already present in the acid whey and formed a specialized homofermentative-lactate-producing microbiome.
- Diluting the acid-whey substrate and shortening the HRT was necessary to alleviate the inhibition effect of the undissociated lactic acid form and to achieve higher lactate conversion rates at high product-into-substrate conversion efficiencies.
- An AFR without a hollow-fiber membrane module and a CSTR with a hollow-fiber membrane module performed better due to a superior cell retention method than a UASB reactor at shorter HRTs.
- Little to no granulation was observed in the UASB reactor. The formed pseudo-granulated biomass resembled a fluffy floc and was quickly lost again.
- During the period between Day 365-384, we experienced an error in the pH control of the CSTR system, where the pH value dropped to 4.3. After this pH-error period, the LG-into-LA conversion efficiency in the CSTR significantly improved and surpassed the AFR.
- A low pH period could be beneficial and lead to increased tolerance of the microbiome to undissociated lactic acid and improve conversion efficiencies.
- The D-lactate-isomer was exclusively produced in all reactor microbiomes that were operated at a pH of 5.0 and temperature of 50°C.

### CRediT authorship contribution statement

Largus T. Angenent (LTA) conceived the project. Richard Hegner (RH), Monika Temovska (MT), and LTA designed the study. MT and RH performed the lab experiments. MT analyzed the data and prepared the figures and tables. MT prepared the 16S rRNA gene amplicon library following the protocol that Andrés E. Ortiz-Ardila (AEO-A) developed. MT performed the bioinformatics and statistical analysis with AEO-A’s advice. The glass reactors were manufactured according to the design by Joseph G. Usack (JGU) and LTA. LTA, RH, AEO-A, and JGU provided guidance. MT and LTA drafted the manuscript. All authors edited the manuscript and approved the final manuscript.

## Acknowledgments

This work was supported by the BMBF VIP + Program (Award # 1336186) and through the Alexander von Humboldt Foundation in the framework of the Alexander von Humboldt Professorship, which was awarded to LTA. This work or part of this work was supported by the Novo Nordisk Foundation CO_2_ Research Center (CORC) with grant number NNF21SA0072700 and is published under the number CORC_24_##. Finally, this work is also supported by the DFG through the Leibniz Prize to LTA. We further acknowledge support by the Max Planck Society to LTA as part of being a Max Planck Fellow. The authors thank Dr. Heike Budde and the Genome Center at the Max Planck Institute (MPI) for Biology Tübingen for performing the MiSeq Illumina sequencing. The authors further wish to thank Dorothea M. Schütterle and Elias Schermuly for helping with the reactor maintenance. In addition, we thank the workers at FrieslandCampina for providing acid whey.

## Supplementary materials

Supplementary data associated with this article can be found, in the online version, at

