## Supplementary Data for "Lactate production from lactose-rich wastewater: A comparative study on reactor configurations to maximize conversion rates and efficiencies"

### S.1 Material and methods

#### *S.1.1 Start-up phase periods*

The start-up phase of the UASB reactor consisted of: **(1)** a 6-day batch mode (Days 0-6); **(2)** a continuous mode at an HRT of 3.7 days, a pH of 5, and a temperature of 44°C (Days 6-10); **(3)** a continuous mode at an HRT of 8.6 days with 2-fold diluted acid whey as the substrate, a pH of 5, and a temperature of 44°C (Days 10-16); **(4)** a continuous mode at an HRT of 1.51 days with 2-fold diluted acid whey as the substrate, a pH of 5.3, and a temperature of 44°C (Days 16-22); and **(5)** a continuous mode at an HRT of 3 days with 2-fold diluted acid whey as the substrate, a pH of 5.3, and a temperature of 44°C (Days 22-35).

The AFR was started up 6 days after the UASB reactor, and the start-up phase consisted of: **(1)** a 10-day batch mode (Days 0-10); **(2)** a continuous mode at an HRT of 1.51 days with 2-fold diluted acid whey as the substrate, a pH of 5.3, and a temperature of 44°C (Days 10-16); and **(3)** a continuous mode at an HRT of 3 days with 2-fold diluted acid whey as the substrate, a pH of 5.3, and a temperature of 44°C (Days 16-29). The CSTR was started up 7 days after the UASB reactor, and the start-up phase consisted of: **(1)** a 9-day batch mode (Days 0-9); **(2)** a continuous mode at an HRT of 1.51 days with 2-fold diluted acid whey as the substrate, a pH of 5.3, and a temperature of 44°C (Days 9-15); and **(3)** a continuous mode at an HRT of 3 days with 2-fold diluted acid whey as the substrate, a pH of 5.3, and a temperature of 44°C (Days 15-28).

#### *S.1.2 PCR cycling conditions*

We performed the 1$^{st}$ PCR using: 2 µL gDNA, 7.5 µL 2X KAPA HiFi HotStart ReadyMix (Roche, Pleasanton CA), 0.6 µL 515F (10 µM) and 0.6 µL 806R (10 µM) primer, 2 µL BSA (10 ppm) solution (T844.4, Carl Roth GmbH, Karlsruhe, Germany), and 2.3 µL PCR-Grade Water (HYCLSH30538.03, VWR International GmbH, Darmstadt, Germany). The PCR cycling conditions were: initial denaturation at 95°C for 3 min, followed by 28 cycles of denaturation at 98°C for 20 s, primer annealing at 67°C for 30 s, extension at 72°C for 20 s, and a final extension at 72°C for 60 s. We performed the 2$^{nd}$ PCR using: 1.5 µL of the 1$^{st}$ PCR product, 7.5 µL 2X KAPA HiFi HotStart ReadyMix, 1.5 µL of each of the Nextera primers for dual indexing, and 3 µL PCR-Grade Water. The PCR cycling conditions were: initial denaturation at 95°C for 3 min, followed by 8 cycles of denaturation at 98°C for 20 s, primer annealing at 70°C for 20 s, extension at 72°C for 45 s, and a final extension at 72°C for 5 min.

### S.2 Equations

#### Hydraulic retention time (HRT) (days):

$$\frac{V}{f}$$

**(Eq. S1)**

Where:

$V$ = reactor wet volume, L

$f$ = effluent flow rate, L $d^{-1}$

#### Volumetric loading rate (mmol C $L^{-1}$ $d^{-1}$):

$$\frac{C_i \times N_C}{HRT}$$

**(Eq. S2)**

Where:

$C_i$ = concentration of influent compound, mmol $L^{-1}$

$N_c$ = conversion factor for mmol to mmol C, which equals the number of carbon atoms of the compound

$HRT$ = hydraulic retention time, d

#### Volumetric SCOD loading rate (g COD $L^{-1}$ $d^{-1}$):

$$\frac{SCOD}{HRT}$$

**(Eq. S3)**

Where:

$SCOD$ = the concentration of soluble chemical oxygen demand in influent, g COD $L^{-1}$

$HRT$ = hydraulic retention time, d

#### Volumetric production rate (mmol C $L^{-1}$ $d^{-1}$):

$$\frac{C_e \times N_C}{HRT}$$

**(Eq. S4)**

Where:

$C_e$ = concentration of effluent compound, mmol $L^{-1}$

$N_c$ = conversion factor for mmol to mmol C, which equals the number of carbon atoms of the compound

$HRT$ = hydraulic retention time, d

### Volumetric conversion rate (mmol C $L^{-1}$ $d^{-1}$):

$$\frac{(C_e \times N_C) - (C_i \times N_C)}{HRT}$$

**(Eq. S5)**

Where:

$C_i$ = concentration of influent compound, mmol $L^{-1}$

$C_e$ = concentration of effluent compound, mmol $L^{-1}$

$N_c$ = conversion factor for mmol to mmol C, which equals the number of carbon atoms of the compound

$HRT$ = hydraulic retention time, d

### LA production specificity (% mmol C):

$$\left(\frac{PR_{LA}}{PR_{LA} + PR_{SP}}\right) * 100$$

**(Eq. S6)**

Where:

$PR_{LA}$ = volumetric lactate production rate, mmol C $L^{-1}$ $d^{-1}$

$PR_{SP}$ = sum of volumetric production rates of the side products ethanol, acetate, and *n*-butyrate, mmol C $L^{-1}$ $d^{-1}$

### LA conversion specificity (% mmol C):

$$\left(\frac{CvR_{LA}}{CvR_{LA} + CvR_{SP}}\right) * 100$$

**(Eq. S7)**

Where:

$CvR_{LA}$ = volumetric lactate conversion rate, mmol C $L^{-1}$ $d^{-1}$

$CvR_{SP}$ = sum of volumetric conversion rates of the side products ethanol, acetate, and *n*-butyrate, mmol C $L^{-1}$ $d^{-1}$

### LG-into-LA conversion efficiency (% mmol C):

$$\left(\frac{CvR_{LA}}{LR_{LG}}\right) * 100$$

**(Eq. S8)**

Where:

$CvR_{LA}$ = volumetric lactate conversion rate, mmol C $L^{-1}$ $d^{-1}$

$LR_{LG}$ = volumetric loading rate of lactose and galactose, mmol C $L^{-1}$ $d^{-1}$

### Volumetric substrate leftover rate (mmol C $L^{-1}$ $d^{-1}$):

$$\frac{(C_{L,e} \times 12) + (C_{G,e} \times 6)}{HRT}$$

**(Eq. S9)**

Where:

$C_{L,e}$ = concentration of lactose in effluent, mmol $L^{-1}$

$C_{G,e}$ = concentration of galactose in effluent, mmol $L^{-1}$

$HRT$ = hydraulic retention time, d

### Volumetric LG consumption rate (mmol C $L^{-1}$ $d^{-1}$):

$$\frac{((C_{L.i} \times 12) - (C_{L,e} \times 12)) + ((C_{G,i} \times 6) - (C_{G,e} \times 6))}{HRT}$$

**(Eq. S10)**

Where:

$C_{L,e}$ = concentration of lactose in effluent, mmol $L^{-1}$

$C_{G,e}$ = concentration of galactose in effluent, mmol $L^{-1}$

$C_{L,i}$ = concentration of lactose in influent, mmol $L^{-1}$

$C_{G,i}$ = concentration of galactose in influent, mmol $L^{-1}$

$HRT$ = hydraulic retention time, d

### LG-into-LA yield (mmol C mmol $C^{-1}$):

$$\left(\frac{CvR_{LA}}{CR_{LG}}\right)$$

**(Eq. S11)**

Where:

$CvR_{LA}$ = volumetric lactate conversion rate, mmol C $L^{-1}$ $d^{-1}$

$CR_{LG}$ = volumetric LG consumption rate, mmol C $L^{-1}$ $d^{-1}$

### LA-to-SCOD effluent concentration ratio (% g COD):

$$\left(\frac{C_{LA,e} \times M_{LA}}{C_{L,e} \times M_L + C_{G,e} \times M_G + C_{LA,e} \times M_{LA} + C_{EtOH,e} \times M_{EtOH} + C_{C2,e} \times M_{C2} + C_{C4,e} \times M_{C4}}\right) * 100$$

**(Eq. S12)**

Where:

$C_{LA,e}$ = lactate concentration in effluent, mmol $L^{-1}$

$C_{L,e}$ = lactose concentration in effluent, mmol $L^{-1}$

$C_{G,e}$ = galactose concentration in effluent, mmol $L^{-1}$

$C_{EtOH,e}$ = ethanol concentration in effluent, mmol $L^{-1}$

$C_{C2,e}$ = acetate concentration in effluent, mmol $L^{-1}$

$C_{C4,e}$ = *n*-butyrate concentration in effluent, mmol $L^{-1}$

$M_{LA}$ = conversion factor for lactate from mmol to g COD, 0.096 g COD $mmol^{-1}$

$M_{L}$ = conversion factor for lactose from mmol to g COD, 0.384 g COD $mmol^{-1}$

$M_{G}$ = conversion factor for galactose from mmol to g COD, 0.192 g COD $mmol^{-1}$

$M_{EtOH}$ = conversion factor for ethanol from mmol to g COD, 0.096 g COD $mmol^{-1}$

$M_{C2}$ = conversion factor for acetate from mmol to g COD, 0.064 g COD $mmol^{-1}$

$M_{C4}$ = conversion factor for *n*-butyrate from mmol to g COD, 0.16 g COD $mmol^{-1}$

### NaOH used for pH regulation (g $L^{-1}$ $d^{-1}$):

$$\frac{\left[\frac{\frac{(W_n - W_{n-1})}{\rho}}{T}\right]}{\frac{1000}{V}} * M * N$$

**(Eq. S13)**

Where:

$W_n$, $W_{n-1}$ = weight of NaOH bottle on day n and n-1, g

$\rho$ = density of 2 M NaOH solution, 1.09 g $cm^3$

$T$ = elapsed time between measurements, d

$V$ = reactor volume, L

$M$ = molar mass of NaOH, 40 g $mol^{-1}$

$N$ = molarity of NaOH solution, 2 M

### S.3 Tables

**Table S1**. Composition of the acid whey batches upon arrival. Reported values are means between all acid whey batches (*n* = 13) used, and the errors represent 95% confidence intervals of mean values. Each acid whey batch was characterized with triplicate measurements except for the solids, COD, and TOC measurements, for which only duplicates were tested.

| **Acid Whey** | |
|---|---|
| Collection site | FrieslandCampina (Cologne, Germany) |
| pH | 4.34 ± 0.02 |
| TS (g $L^{-1}$) | 61.0 ± 1.21 |
| VS (g $L^{-1}$) | 53.4 ± 1.16 |
| TCOD (g COD $L^{-1}$) | 69.6 ± 1.65 |
| SCOD (g COD $L^{-1}$) | 65.7 ± 1.63 |
| TOC (g C $L^{-1}$) | 22.9 ± 0.34 |
| DOC (g C $L^{-1}$) | 22.7 ± 0.33 |
| SCOD-to-DOC ratio | 2.90 ± 0.11 |
| Lactose (mmol $L^{-1}$) | 110 ± 1.46 |
| Galactose (mmol $L^{-1}$) | 36.4 ± 0.63 |
| Lactate (mmol $L^{-1}$) | 103 ± 5.24 |
| Lactose (mmol C $L^{-1}$) | 1323 ± 17.6 |
| Galactose (mmol C $L^{-1}$) | 218 ± 3.81 |
| Lactate (mmol C $L^{-1}$) | 310 ± 15.7 |
| Lactose* (g COD $L^{-1}$) | 42.3 ± 0.56 |
| Galactose* (g COD $L^{-1}$) | 6.99 ± 0.12 |
| Lactate* (g COD $L^{-1}$) | 9.92 ± 0.50 |
| Lactose** (g C $L^{-1}$) | 15.9 ± 0.21 |
| Galactose** (g C $L^{-1}$) | 2.62 ± 0.05 |
| Lactate** (g C $L^{-1}$) | 3.72 ± 0.19 |
| Lactose-to-Lactate ratio (mmol C $L^{-1}$) | 4.37 ± 0.22 |
| Calculated SCOD* (g COD $L^{-1}$) | 59.2 ± 0.88 |
| Calculated SCOD*-to-SCOD ratio | 0.90 ± 0.03 |
| Calculated DOC** (g C $L^{-1}$) | 22.2 ± 0.33 |
| Calculated DOC**-to-DOC ratio | 0.98 ± 0.02 |
| Calculated SCOD*-to-DOC** ratio | 2.66 |

* COD conversions are 0.384, 0.192, and 0.096 g COD $mmol^{-1}$ for lactose, galactose, and lactate, respectively.
** DOC conversions are 0.144, 0.072, and 0.036 g C $mmol^{-1}$ for lactose, galactose, and lactate, respectively. The total calculated SCOD and DOC include lactose, galactose, and lactate, the predominant sources of SCOD and DOC in acid whey.

**Table S2.** The operating and performance parameters for the UASB reactor. Reported values are means between: **(1)** 25; **(2)** 16; **(3)** 15; and **(4)** 15 days of steady-state operation at the given HRT of 3, 2, 1, and 0.6 days, respectively. Errors represent 95% confidence intervals of mean values.

| | **UASB reactor** | | | |
|---|---|---|---|---|
| **HRT (d)** | **3.04** | **2.16** | **1.00** | **0.61** |
| Volumetric LGLA loading rate (mmol C $L^{-1}$ $d^{-1}$) | 600 ± 8.85 | 760 ± 30.34 | 1759 ± 34.9 | 1693 ± 22.5 |
| Volumetric LG loading rate (mmol C $L^{-1}$ $d^{-1}$) | 492 ± 9.48 | 620 ± 25.10 | 1439 ± 28.6 | 1446 ± 17.0 |
| Volumetric lactose loading rate (mmol C $L^{-1}$ $d^{-1}$) | 446 ± 4.78 | 539 ± 21.7 | 1245 ± 23.7 | 1273 ± 15.2 |
| Volumetric galactose loading rate (mmol C $L^{-1}$ $d^{-1}$) | 45.3 ± 6.93 | 81.3 ± 3.84 | 194 ± 9.39 | 173 ± 2.71 |
| Volumetric EtOH loading rate (mmol C $L^{-1}$ $d^{-1}$) | 15.4 ± 4.38 | 0.60 ± 0.68 | 7.30 ± 3.14 | 0.00 |
| Volumetric C2 loading rate (mmol C $L^{-1}$ $d^{-1}$) | 0.89 ± 0.20 | 0.00 | 0.22 ± 0.32 | 0.00 |
| Volumetric C4 loading rate (mmol C $L^{-1}$ $d^{-1}$) | 0.00 | 0.00 | 0.00 | 0.00 |
| Volumetric SCOD loading rate (g COD $L^{-1}$ $d^{-1}$) | 20.0 ± 0.19 | 24.3 ± 0.98 | 56.6 ± 1.10 | 54.2 ± 0.72 |
| Wet volume (L) | 2.8 | | | |
| Flow rate (L $d^{-1}$) | 0.92 | 1.30 | 2.79 | 4.61 |
| Upflow velocity (m $h^{-1}$) | 3.92 | 3.92 | 3.94 | 3.97 |
| NaOH used for pH regulation (g $L^{-1}$ $d^{-1}$) | 1.92 ± 0.25 | 4.08 ± 0.55 | 6.39 ± 0.67 | 10.2 ± 0.73 |
| % Acid whey influent (times diluted) | 100 (0) | 100 (0) | 100 (0) | 60 (1.67) |
| Volumetric lactose consumption rate (mmol C $L^{-1}$ $d^{-1}$) | 163 ± 15.0 | 200 ± 20.1 | 588 ± 35.3 | 726 ± 49.7 |
| Volumetric galactose consumption rate (mmol C $L^{-1}$ $d^{-1}$) | 43.1 ± 6.94 | 78.6 ± 3.59 | 151 ± 14.2 | 158 ± 5.03 |
| Volumetric LA production rate (mmol C $L^{-1}$ $d^{-1}$) | 267 ± 6.91 | 441 ± 9.18 | 1066 ± 51.0 | 1059 ± 45.6 |
| Volumetric LA conversion rate (minus LA in influent) (mmol C $L^{-1}$ $d^{-1}$) | 158. ± 6.72 | 297 ± 9.87 | 747 ± 45.8 | 812 ± 45.1 |
| LA production specificity (% mmol C) | 92.8 ± 0.64 | 99.7 ± 0.21 | 99.5 ± 0.13 | 100 ± 0.03 |
| LA conversion specificity (% mmol C) | 95.4 ± 1.42 | 99.7 ± 0.25 | 99.6 ± 0.04 | 100 ± 0.04 |
| LG-into-LA conversion efficiency (% mmol C) | 32.8 ± 1.38 | 47.2 ± 2.16 | 51.9 ± 3.17 | 56.1 ± 2.98 |
| LG-into-LA yield (mmol C mmol $C^{-1}$) | 0.78 ± 0.04 | 1.04 ± 0.04 | 1.01 ± 0.04 | 0.92 ± 0.02 |
| LA-to-SCOD effluent conc. ratio (% g COD) | 45.9 ± 1.43 | 56.3 ± 1.46 | 60.1 ± 2.26 | 65.3 ± 2.80 |
| Undissociated LA (mmol C $L^{-1}$) | 55.3 ± 1.94 | 64.3 ± 1.34 | 72.2 ± 3.45 | 45.9 ± 4.87 |
| Experimental $gNaOH_{used}\_g^{-1}\_LA_{converted}$ | 0.40 ± 0.05 | 0.46 ± 0.07 | 0.30 ± 0.05 | 0.42 ± 0.02 |
| Theoretical $gNaOH_{used}\_g^{-1}\_LA_{converted}$ | 0.44 | | | |
| Volumetric EtOH production rate (mmol C $L^{-1}$ $d^{-1}$) | 17.3 ± 2.17 | 0.77 ± 0.84 | 2.49 ± 1.43 | 0.00 |
| Volumetric EtOH conversion rate (minus EtOH in influent) (mmol C $L^{-1}$ $d^{-1}$) | 5.26 ± 2.54 | 0.27 ± 0.45 | 0.00 | 0.00 |
| Volumetric C2 production rate (mmol C $L^{-1}$ $d^{-1}$) | 2.47 ± 0.07 | 0.57 ± 0.39 | 3.35 ± 0.39 | 0.28 ± 0.37 |
| Volumetric C2 conversion rate (minus C2 in influent) (mmol C $L^{-1}$ $d^{-1}$) | 1.59 ± 0.19 | 0.54 ± 0.38 | 3.13 ± 0.37 | 0.28 ± 0.37 |
| Volumetric C4 production rate (mmol C $L^{-1}$ $d^{-1}$) | 1.14 ± 0.29 | 0.00 | 0.00 | 0.00 |
| Volumetric C4 conversion rate (minus C4 in influent) (mmol C $L^{-1}$ $d^{-1}$) | 1.14 ± 0.29 | 0.00 | 0.00 | 0.00 |
| Side products (EtOH, C2, C4) yield (mmol C mmol $C^{-1}$) | 0.04 ± 0.01 | $3.28x10^{-3}$ ± $2.58x10^{-3}$ | $4.24x10^{-3}$ ± $4.29x10^{-4}$ | $3.13x10^{-4}$ ± $4.18x10^{-4}$ |

**Table S3.** The operating and performance parameters for the AFR. Reported values are means between: **(1)** 25; **(2)** 16; **(3)** 15; and **(4)** 15 days of steady-state operation at the given HRT of 3, 2, 1, and 0.6 days, respectively. Errors represent 95% confidence intervals of mean values.

| | **AFR** | | | |
|---|---|---|---|---|
| **HRT (d)** | **3.06** | **2.18** | **1.02** | **0.61** |
| Volumetric LGLA loading rate (mmol C $L^{-1}$ $d^{-1}$) | 586 ± 7.32 | 640 ± 92.3 | 1692 ± 47.2 | 1671 ± 27.9 |
| Volumetric LG loading rate (mmol C $L^{-1}$ $d^{-1}$) | 479 ± 7.74 | 622 ± 12.2 | 1377 ± 35.8 | 1420 ± 24.8 |
| Volumetric lactose loading rate (mmol C $L^{-1}$ $d^{-1}$) | 443 ± 2.59 | 541 ± 10.1 | 1191 ± 31.6 | 1242 ± 21.6 |
| Volumetric galactose loading rate (mmol C $L^{-1}$ $d^{-1}$) | 35.8 ± 7.45 | 81.0 ± 2.78 | 186 ± 9.94 | 177 ± 6.11 |
| Volumetric EtOH loading rate (mmol C $L^{-1}$ $d^{-1}$) | 19.6 ± 4.33 | 0.29 ± 0.40 | 4.28 ± 2.87 | 0.00 |
| Volumetric C2 loading rate (mmol C $L^{-1}$ $d^{-1}$) | 0.88 ± 0.11 | 0.00 | 0.21 ± 0.31 | 0.00 |
| Volumetric C4 loading rate (mmol C $L^{-1}$ $d^{-1}$) | 0.00 | 0.00 | 0.00 | 0.00 |
| Volumetric SCOD loading rate (g COD $L^{-1}$ $d^{-1}$) | 19.7 ± 0.10 | 24.0 ± 0.94 | 54.3 ± 1.55 | 53.5 ± 0.89 |
| Wet volume (L) | 2.2 | | | |
| Flow rate (L $d^{-1}$) | 0.72 | 1.01 | 2.16 | 3.60 |
| Upflow velocity (m $h^{-1}$) | 1.92 | 1.92 | 1.94 | 1.96 |
| NaOH used for pH regulation (g $L^{-1}$ $d^{-1}$) | 2.26 ± 0.25 | 3.57 ± 0.32 | 6.27 ± 0.42 | 10.9 ± 0.62 |
| % Acid whey influent (times diluted) | 100 (0) | 100 (0) | 100 (0) | 60 (1.67) |
| Volumetric lactose consumption rate (mmol C $L^{-1}$ $d^{-1}$) | 208 ± 14.5 | 366 ± 13.6 | 619 ± 26.3 | 993 ± 27.7 |
| Volumetric galactose consumption rate (mmol C $L^{-1}$ $d^{-1}$) | 29.1 ± 7.51 | 20.7 ± 9.32 | 0.00 | 71.5 ± 12.6 |
| Volumetric LA production rate (mmol C $L^{-1}$ $d^{-1}$) | 293 ± 10.3 | 519 ± 13.8 | 886 ± 16.1 | 1194 ± 22.1 |
| Volumetric LA conversion rate (minus LA in influent) (mmol C $L^{-1}$ $d^{-1}$) | 186 ± 10.4 | 380 ± 14.1 | 572 ± 17.5 | 942 ± 24.6 |
| LG-into-LA conversion efficiency (% mmol C) | 38.9 ± 2.23 | 61.2 ± 2.64 | 41.7 ± 1.91 | 66.5 ± 2.17 |
| LA production specificity (% mmol C) | 91.0 ± 0.83 | 97.8 ± 0.34 | 98.7 ± 0.43 | 96.7 ± 0.43 |
| LA conversion specificity (% mmol C) | 94.1 ± 1.37 | 91.4 ± 11.6 | 98.3 ± 0.55 | 95.9 ± 0.56 |
| LG-into-LA yield (mmol C mmol $C^{-1}$) | 0.79 ± 0.03 | 0.98 ± 0.03 | 0.93 ± 0.05 | 0.89 ± 0.03 |
| LA-to-SCOD effluent conc. ratio (% g COD) | 51.0 ± 2.09 | 67.3 ± 1.75 | 52.5 ± 1.30 | 75.3 ± 1.55 |
| Undissociated LA (mmol C $L^{-1}$) | 60.0 ± 2.00 | 76.5 ± 2.03 | 60.9 ± 1.11 | 49.3 ± 0.91 |
| Experimental $gNaOH_{used}\_g^{-1}\_LA_{converted}$ | 0.40 ± 0.03 | 0.31 ± 0.03 | 0.37 ± 0.02 | 0.39 ± 0.02 |
| Theoretical $gNaOH_{used}\_g^{-1}\_LA_{converted}$ | 0.44 | | | |
| Volumetric EtOH production rate (mmol C $L^{-1}$ $d^{-1}$) | 20.7 ± 2.19 | 4.11 ± 0.31 | 1.32 ± 1.31 | 1.48 ± 1.55 |
| Volumetric EtOH conversion rate (minus EtOH in influent) (mmol C $L^{-1}$ $d^{-1}$) | 4.83 ± 2.56 | 3.82 ± 0.59 | 0.03 ± 0.06 | 1.48 ± 1.55 |
| Volumetric C2 production rate (mmol C $L^{-1}$ $d^{-1}$) | 3.26 ± 0.14 | 2.44 ± 0.49 | 3.69 ± 1.20 | 7.98 ± 1.31 |
| Volumetric C2 conversion rate (minus C2 in influent) (mmol C $L^{-1}$ $d^{-1}$) | 2.37 ± 0.14 | 2.44 ± 0.49 | 3.48 ± 1.13 | 7.98 ± 1.31 |
| Volumetric C4 production rate (mmol C $L^{-1}$ $d^{-1}$) | 4.65 ± 0.83 | 4.95 ± 1.17 | 6.61 ± 2.10 | 31.1 ± 4.17 |
| Volumetric C4 conversion rate (minus C4 in influent) (mmol C $L^{-1}$ $d^{-1}$) | 4.65 ± 0.83 | 4.95 ± 1.17 | 6.61 ± 2.10 | 31.1 ± 4.17 |
| Side products (EtOH, C2, C4) yield (mmol C mmol $C^{-1}$) | 0.05 ± 0.01 | 0.03 ± 0.01 | $0.02 \pm 4.87 \times 10^{-3}$ | $0.04 \pm 4.51 \times 10^{-3}$ |

**Table S4.** The operating and performance parameters for the CSTR. Reported values are means between: **(1)** 25; **(2)** 16; **(3)** 15; and **(4)** 15 days of steady-state operation at the given HRT of 3, 2, 1, and 0.6 days, respectively. The 0.60-I-day HRT represents data before the pH-error period. The 0.60-II-day HRT represents data after the pH-error period. Errors represent 95% confidence intervals of mean values.

| | | | **CSTR** | | |
|---|---|---|---|---|---|
| **HRT (d)** | **2.99** | **1.99** | **1.00** | **0.60-I** | **0.60-II** |
| Volumetric LGLA loading rate (mmol C $L^{-1}$ $d^{-1}$) | 600 ± 7.59 | 837 ± 18.1 | 1740 ± 41.7 | 1719 ± 42.9 | 1800 ± 39.7 |
| Volumetric LG loading rate (mmol C $L^{-1}$ $d^{-1}$) | 490 ± 8.00 | 683 ± 14.7 | 1424 ± 35.9 | 1383 ± 36.1 | 1531 ± 34.7 |
| Volumetric lactose loading rate (mmol C $L^{-1}$ $d^{-1}$) | 453 ± 2.58 | 594 ± 12.0 | 1252 ± 27.7 | 1200 ± 30.9 | 1345 ± 30.5 |
| Volumetric galactose loading rate (mmol C $L^{-1}$ $d^{-1}$) | 36.7 ± 7.59 | 89.5 ± 3.29 | 172 ± 14.6 | 182 ± 7.32 | 186 ± 8.44 |
| Volumetric EtOH loading rate (mmol C $L^{-1}$ $d^{-1}$) | 19.9 ± 4.37 | 0.69 ± 0.76 | 10.7 ± 5.38 | 0.00 | 0.00 |
| Volumetric C2 loading rate (mmol C $L^{-1}$ $d^{-1}$) | 0.91 ± 0.11 | 0.00 | 0.00 | 0.00 | 0.00 |
| Volumetric C4 loading rate (mmol C $L^{-1}$ $d^{-1}$) | 0.00 | 0.00 | 0.00 | 0.00 | 0.00 |
| Volumetric SCOD loading rate (g COD $L^{-1}$ $d^{-1}$) | 20.2 ± 0.10 | 26.8 ± 0.59 | 56.2 ± 1.20 | 55.0 ± 1.37 | 57.6 ± 1.27 |
| Wet volume (L) | | | 1.72 | | |
| Flow rate (L $d^{-1}$) | 0.58 | 0.86 | 1.73 | 2.88 | 2.88 |
| NaOH used for pH regulation (g $L^{-1}$ $d^{-1}$) | 2.02 ± 0.36 | 3.09 ± 0.38 | 8.22 ± 1.05 | 10.1 ± 0.22 | 17.3 ± 0.88 |
| % Acid whey influent (times diluted) | 100 (0) | 100 (0) | 100 (0) | 60 (1.67) | 60 (1.67) |
| Volumetric lactose consumption rate (mmol C $L^{-1}$ $d^{-1}$) | 142 ± 14.6 | 357 ± 22.3 | 543 ± 48.0 | 748 ± 34.5 | 1310 ± 45.4 |
| Volumetric galactose consumption rate (mmol C $L^{-1}$ $d^{-1}$) | 31.6 ± 7.82 | 84.3 ± 4.21 | 157 ± 13.6 | 135 ± 12.7 | 174 ± 9.86 |
| Volumetric LA production rate (mmol C $L^{-1}$ $d^{-1}$) | 246 ± 10.9 | 561 ± 21.2 | 979 ± 47.7 | 1153 ± 17.0 | 1525 ± 41.8 |
| Volumetric LA conversion rate (minus LA in influent) (mmol C $L^{-1}$ $d^{-1}$) | 136 ± 11.0 | 408 ± 19.7 | 663 ± 40.3 | 817 ± 19.8 | 1256 ± 46.3 |
| LA production specificity (% mmol C) | 90.0 ± 0.94 | 92.8 ± 2.98 | 99.8 ± 0.09 | 99.7 ± 0.24 | 99.8 ± 0.16 |
| LA conversion specificity (% mmol C) | 93.9 ± 2.25 | 97.8 ± 0.28 | 99.7 ± 0.13 | 99.6 ± 0.34 | 99.8 ± 0.21 |
| LG-into-LA conversion efficiency (% mmol C) | 27.9 ± 2.38 | 59.8 ± 3.17 | 46.5 ± 2.03 | 59.2 ± 1.80 | 82.2 ± 3.41 |
| LG-into-LA yield (mmol C mmol $C^{-1}$) | 0.79 ± 0.03 | 0.93 ± 0.04 | 0.96 ± 0.05 | 0.93 ± 0.04 | 0.85 ± 0.02 |
| LA-to-SCOD effluent conc. ratio (% g COD) | 40.9 ± 2.12 | 68.9 ± 2.21 | 57.3 ± 2.00 | 69.6 ± 1.01 | 96.9 ± 2.37 |
| Undissociated LA (mmol C $L^{-1}$) | 49.6 ± 2.20 | 75.5 ± 2.85 | 65.8 ± 3.21 | 46.5 ± 0.68 | 62.5 ± 2.84 |
| Experimental $gNaOH_{used}\_g^{-1}\_LA_{converted}$ | 0.49 ± 0.07 | 0.26 ± 0.03 | 0.41 ± 0.05 | 0.41 ± 0.01 | 0.46 ± 0.01 |
| Theoretical $gNaOH_{used}\_g^{-1}\_LA_{converted}$ | | | 0.44 | | |
| Volumetric EtOH production rate (mmol C $L^{-1}$ $d^{-1}$) | 24.1 ± 2.60 | 3.43 ± 0.93 | 0.00 | 3.24 ± 2.73 | 0.00 |
| Volumetric EtOH conversion rate (minus EtOH in influent) (mmol C $L^{-1}$ $d^{-1}$) | 6.80 ± 3.28 | 2.88 ± 1.00 | 0.00 | 3.24 ± 2.73 | 0.00 |
| Volumetric C2 production rate (mmol C $L^{-1}$ $d^{-1}$) | 3.10 ± 0.19 | 2.97 ± 0.46 | 2.25 ± 0.84 | 0.00 | 2.54 ± 2.23 |
| Volumetric C2 conversion rate (minus C2 in influent) (mmol C $L^{-1}$ $d^{-1}$) | 2.19 ± 0.23 | 2.97 ± 0.46 | 2.25 ± 0.84 | 0.00 | 2.54 ± 2.23 |
| Volumetric C4 production rate (mmol C $L^{-1}$ $d^{-1}$) | 0.00 | 3.19 ± 1.07 | 0.00 | 0.00 | 0.00 |
| Volumetric C4 conversion rate (minus C4 in influent) (mmol C $L^{-1}$ $d^{-1}$) | 0.00 | 3.19 ± 1.07 | 0.00 | 0.00 | 0.00 |
| Side products (EtOH, C2, C4) yield (mmol C mmol $C^{-1}$) | 0.05 ± 0.02 | 0.02 ± $2.79 \times 10^{-3}$ | $3.20 \times 10^{-3}$ ± $1.20 \times 10^{-3}$ | $3.69 \times 10^{-3}$ ± $3.22 \times 10^{-3}$ | $1.80 \times 10^{-3}$ ± $1.66 \times 10^{-3}$ |

### S.4 Figures

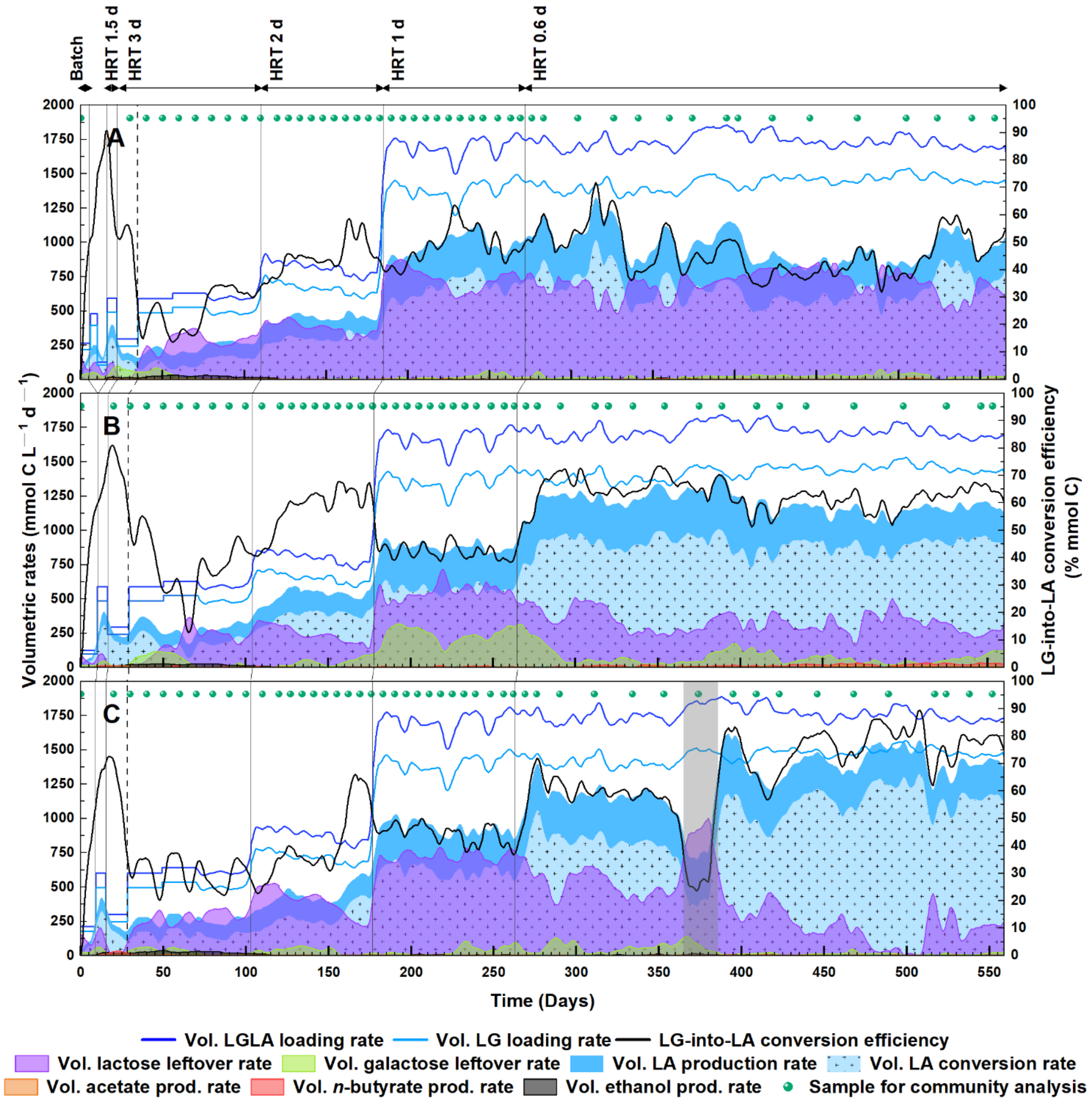


**Fig. S1.** Performance of the reactor microbiomes during the total operating period. (A) UASB reactor; (B) AFR; and (C) CSTR. Area plots of the lactate production rate, lactate conversion rate, side products (ethanol, acetate, and *n*-butyrate) production rates, and leftover substrate rates (lactose and galactose) on the left axes. Line plots of the total organic loading rate (lactose, galactose, and lactate) and the substrate loading rate (lactose and galactose) on the left axes. Line plots of the LG-into-LA conversion efficiencies on the right axes. The data represent a 6-day moving average (each average contains six sampling points). Dash lines mark the end of the start-up phase and note the start of the comparison periods. The time period for the CSTR marked with gray represents the pH-error period when the actual pH value was 4.3.Vol. in volumetric.

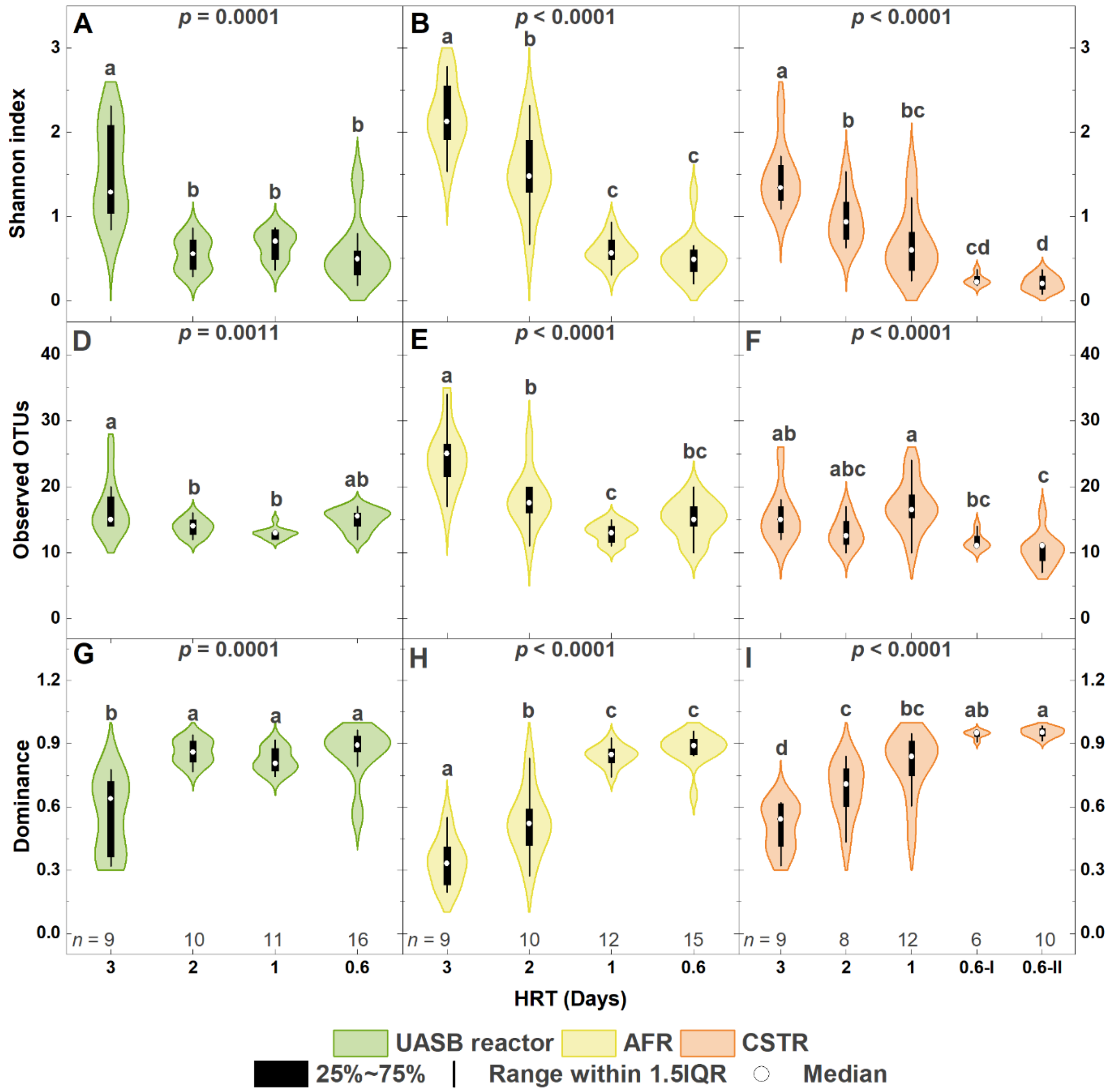


**Fig. S2.** Comparisons of α-diversity metrics for the individual reactor microbiomes\` during the different operating HRTs. (A-C) Shannon Index; (D-E) observed OTUs (operational taxonomic unit); and (G-E) dominance. Significant differences based on the Kruskal-Wallis test ($p$ < 0.05) between the reactor microbiomes at a given HRT are indicated with significance letters. For means denoted with the same letter, the difference between the means is not statistically significant.

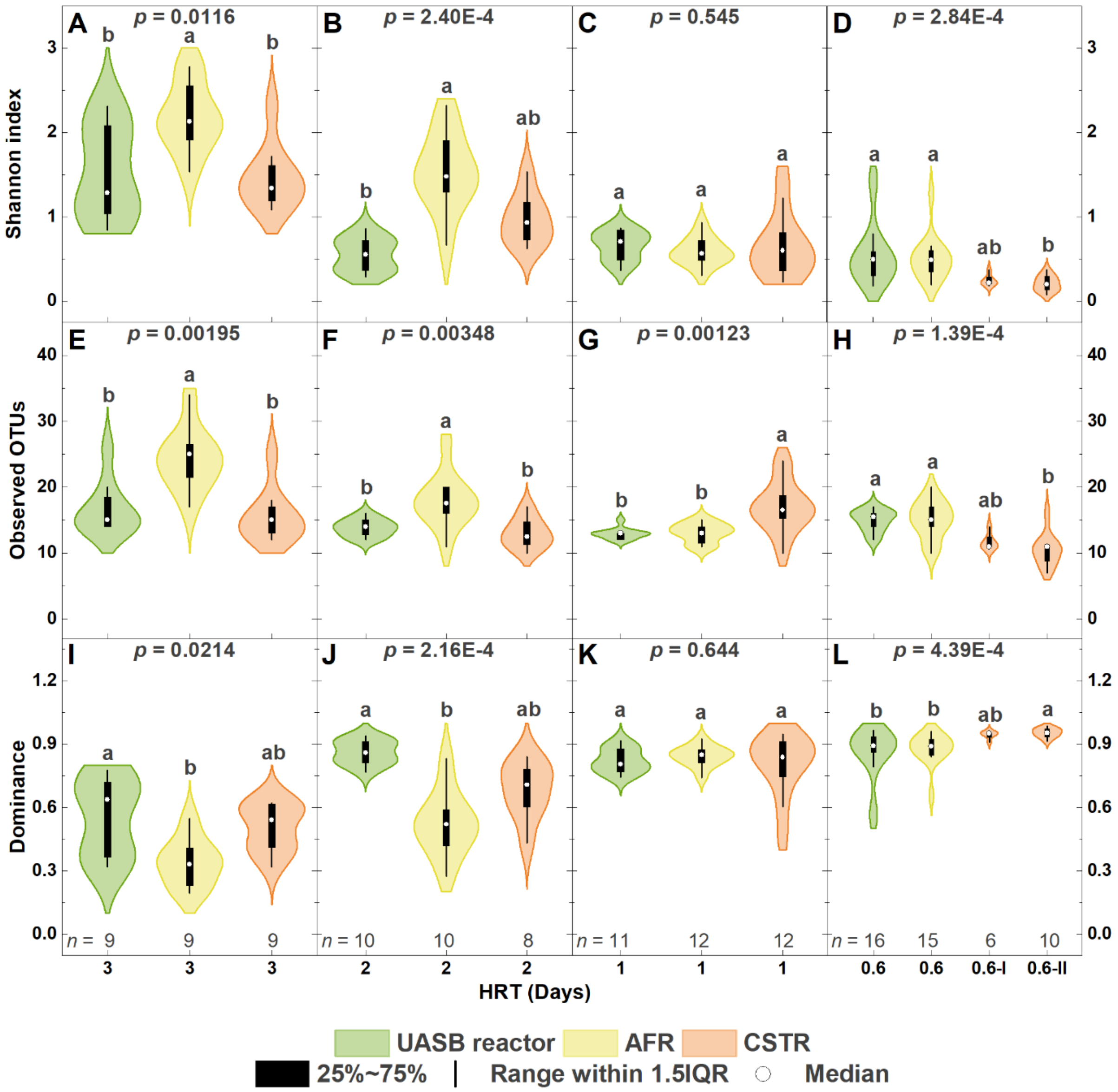


**Fig. S3.** Comparisons of α-diversity metrics between the three reactor microbiomes` during the different operating HRTs. **(**A-D) Shannon Index; (E-H) observed OTUs (operational taxonomic unit); and (I-L) dominance. Significant differences based on the Kruskal-Wallis test ($p$ < 0.05) between the reactor microbiomes at a given HRT are indicated with significance letters. For means denoted with the same letter, the difference between the means is not statistically significant.
